# Comparison of Four Assays That Measure Antibodies to Ebola Virus Glycoprotein

**DOI:** 10.64898/2026.03.18.708022

**Authors:** Irina Maljkovic Berry, Safaa Ben Farhat, Viviane Callier, Céline Roy, Natasha Dubois Cauwelaert, Edouard Lhomme, Prabha Chandrasekaran, Amie Jarra, Harrison Gichini, Scott Anthony, Nicolas Bernaud, Christine Schwimmer, Martine Peeters, Guillaume Thaurignac, Neraide Biai, Stephen B. Kennedy, Mark Kieh, Sarah M. Browne, Mosoka Fallah, Patrick Mutombo, Emmanuel Lokilo, Olivier Tshiani Mbaya, Lisa Hensley, Ian Crozier, Richard T. Davey, Yves Lévy, Ahidjo Ayouba, Laura Richert, H. Clifford Lane, Cavan Reilly, Dean A. Follmann

**Affiliations:** Integrated Research Facility at Fort Detrick, National Institute of Allergy and Infectious Diseases, National Institutes of Health, Fort Detrick, Frederick, MD 21702, USA; University of Bordeaux, INSERM, Institut Bergonié, CHU de Bordeaux, CIC 1401, Euclid/F-CRIN clinical trials platform, F-33000 Bordeaux, France; Clinical Monitoring Research Program Directorate, Frederick National Laboratory for Cancer Research, Frederick, MD 21702, USA; University of Bordeaux, INSERM, MART, UMS 54, F-33000 Bordeaux, France; French Institute for Health and Medical Research (INSERM), 75013 Paris, France; ANRS Emerging Infectious Diseases, Paris, France; Vaccine Research Institute, Univ. Paris Est Créteil, Henri Mondor Hospital, Créteil, France; Recherche Translationnelle Appliquée au VIH et aux Maladies Infectieuses, University of Montpellier, Institut de Recherche pour le Développement, INSERM, 34090, Montpellier, France; Partnership for Research on Vaccines and Infectious Diseases in Liberia (PREVAIL), Monrovia, 1000 Liberia; National Public Health Institute of Liberia, Monrovia, Liberia; Laboratory of Immunoregulation, National Institute of Allergy and Infectious Diseases, National Institutes of Health, Bethesda, MD 20892, USA; Institut de Recherche Biomédicale (INRB), Kinshasa – Gombe, DRC; U1219 BPH, Inria, Sistm, F-33000 Bordeaux, France; Division of Biostatistics and Health Data Science, University of Minnesota, Minneapolis, MN, USA; Biostatistics Research Branch, National Institute of Allergy and Infectious Diseases, National Institutes of Health, Bethesda, MD 20892, USA

**Keywords:** Ebola virus, glycoprotein, ELISA, assay comparison, antibody

## Abstract

The accurate measurement of Ebola virus (EBOV)-specific antibody responses is crucial to assessing immunity induced by EBOV infection or vaccination. For this purpose, the Filovirus Animal Nonclinical Group (FANG) anti-EBOV glycoprotein (GP_1,2_) ELISA is considered the “gold-standard”. However, it has limitations such as high repeat-rates and variability, and low throughput. Here, we describe two new alternative assays: a Single-Molecule Assay Planar EBOV GP_1,2_ ELISA and a multiplexed EBOV GP_1,2_, EBOV nucleoprotein, and EBOV Viral Protein 40 Luminex assay, and compare these with two versions of the FANG ELISA. Samples were selected from participants receiving vaccine or placebo in a randomized, placebo-controlled, double-blinded study of two EBOV vaccines (PREVAIL 1), and a longitudinal cohort study of Ebola virus disease (EVD) survivors and their close contacts (PREVAIL 3). All four assays were concordant in their measurements of anti-EBOV GP_1,2_-specific immunoglobulin G responses, allowing for the determination of conversion equations for antibody measurements across assays. In addition, all four showed a similar ability to distinguish vaccine recipients from placebo recipients and EVD survivors from their close contacts. Compared to the FANG assays, the Quanterix and Luminex assays had lower variability, lower repeat rates, and higher throughput, making them good alternatives for future studies.

## INTRODUCTION

Before 2013, outbreaks of Ebola virus disease (EVD) occurred primarily in rural areas of Middle Africa and were sporadic and limited in scale and scope. In contrast, the 2013–2016 Western African EVD outbreak involved multiple countries and major urban areas and resulted in approximately 28,000 infections and 11,000 deaths (1, 2). The rapid spread, scope, and scale of the outbreak prompted the declaration of a Public Health Emergency of International Concern by the World Health Organization (WHO) and marked acceleration of countermeasure development and evaluation (3, 4). In the affected region, several clinical trials were quickly established to identify safe and effective vaccines to prevent Ebola virus (EBOV) infection and EVD. A cluster-randomized ring trial of the rVSVΔG-ZEBOV-GP (rVSV-ZEBOV) vaccine employing immediate versus deferred immunizations of contacts reported 100% efficacy when excluding infections within the first nine days (95% confidence interval: 79.3%, 100%) (4, 5). As the epidemic waned in 2016, evaluating vaccine efficacy became challenging. Thus, the focus of ongoing clinical trials shifted towards evaluating the safety and immunogenicity, specifically in the measurement of anti-EBOV glycoprotein (GP_1,2_) immunoglobulin G (IgG) levels previously identified in animal models as potential immune correlates of vaccine-mediated protection (6).

Prior to the 2013–2016 outbreak, anti-EBOV-GP IgG assays had been developed for small-scale research use and were not optimized for field evaluation of medical countermeasures. In the absence of sufficient characterization, standardization, and validation, considerable uncertainty around assay accuracy, consistency, and reproducibility limited their usefulness in the regulatory context, such as for vaccine approval. To address this gap, the Filovirus Animal Nonclinical Group (FANG) developed and subsequently validated an anti-EBOV GP_1,2_ IgG enzyme-linked immunosorbent assay (ELISA) for the detection and quantitation of EBOV-GP_1,2_-specific IgG in human serum (7). The FANG assay had greater precision, lower background, and a broader range of accuracy compared to the commercially available Alpha Diagnostic International assay (8). The FANG assay has since been established as the “gold standard” for measuring humoral responses following EBOV vaccination or after EVD (8–12).

However, challenges remain with the FANG assay, including low throughput due to the need for numerous sample dilution series, frequent sample or plate failures that require repeat testing, and samples often falling outside the assay dynamic range, necessitating additional dilutions.

Furthermore, substantial inter-laboratory variability in FANG assay results has prompted some studies to use a single laboratory only for testing (12–14). Additionally, though both have been used in the research setting, the qualified “old” FANG (oFANG) and the validated “new” FANG (nFANG) versions of the assay differ in reference standard dilutions, assay limit of detection (LOD), lower limit of quantitation (LLOQ), upper limit of quantitation (ULOQ), assay sample passing algorithms, and in how sample replicates are assayed across plates, making direct comparisons challenging. These challenges made the FANG assay less suitable for comparisons between laboratories or studies, or for studies with large sample numbers or samples collected over long time periods. The limitations of the two FANG assays prompted the development of other EBOV antigen-specific assays of IgG (15–17), including the two assays evaluated in this report.

The quantitative Single-Molecule Assay (SIMOA) Planar Ebola GP_1,2_ ELISA (Q2) on the Quanterix SP-X platform is a multiplex-based sandwich ELISA in a planar, microplate-based array format in which each well is pre-spotted with EBOV GP_1,2_ monomer proteins in a circular pattern. SIMOA planar digital technology enables higher sensitivity compared to standard ELISAs, achieving LODs in attomolar range (10^-18^ compared to standard 10^-9^ to 10^-12^) (18, 19). These assays have been shown to have equal or better sensitivity than other ELISAs for antibody measurements, as well as a larger dynamic range (20). The second assay, the Luminex-based multiplex immunoassay, enables the simultaneous detection of antibodies in response to multiple EBOV antigens (16). The Luminex assay utilizes carboxyl-functionalized fluorescent magnetic beads coupled with recombinant EBOV antigens, including nucleoprotein (NP), viral protein 40 (VP40), and GP_1,2_ from different EBOV isolates, specifically those from the 1976 Yambuku (Yambukua-Mayinga variant/isolate) and 1995 Kikwit (Kikwit/9510621 variant/isolate) EVD outbreaks. In general, Luminex bead-based assays offer several advantages over traditional ELISAs, including higher throughput, lower sample volume requirements, and the ability to simultaneously measure responses to multiple antigens (21, 22).

The use of different assays to measure EBOV GP_1,2_ antibodies limits the ability to compare and synthesize results, and to establish standardized correlates of protection—ultimately limiting progress in vaccine evaluation and regulatory decision making. Additionally, the lack of head-to-head comparison prevents the accurate assessment of the assays’ relative performance characteristics. Here, we compare the performance of four assays (Q2, Luminex, oFANG, and nFANG) by analyzing antibody responses in samples from rVSV-ZEBOV vaccine and placebo recipients, and in EVD survivors and their close contacts. In addition, we provide equations for converting one assay readout to another. We also evaluate the assays on a range of practical characteristics relevant to their use in a field laboratory.

## RESULTS

### Participants

A total of 149 samples were selected from the Partnership for Research on Vaccines and Infectious Diseases in Liberia (PREVAIL) 1 study, including 50 from the placebo arm and 99 from the vaccine arms (Table S1.1). In this population, the mean age was 34 years, and 29% of participants were female. A total of 149 samples were selected from PREVAIL 3, including 120 EVD survivors (mean age 34 years, 55% female) and 29 close contacts (mean age 28 years, 59% female).

### Assay Validation

Q2 and Luminex assay performance in specifically detecting anti-EBOV GP_1,2_ IgG, including sensitivity, specificity, and accuracy; the upper and lower limits of assay quantitation, and intra-and inter-assay variability was assessed through assay validation (Supplemental Material). Both assays were shown to be specific and passed pre-determined assay validation criteria. Intra-assay variability based on high, low, and reference standard quality controls was slightly lower for the Q2 assay, with an average coefficient of variation (CV) of 6.7% (range: 5.1%–9.1%), compared to 12.4% for the Luminex assay (range: 11.72%–13.51%). Likewise, inter-assay variability was lower for the Q2 assay, with an average CV of 10.7% (range 6.0%–24.7%), compared to the Luminex average CV of 17.2% (range 14.03%–20.80%).

### Assay Concordance and Performance

A comparison of anti-EBOV GP_1,2_ IgG concentrations ([IgG]) measured in 298 samples across all four assays (oFANG, nFANG, Q2, and Luminex) showed similar assay readouts across the four study groups, with the Q2 assay [IgG] being somewhat lower in the EVD survivor group compared to the other assays, and the oFANG assay [IgG] higher in both the placebo and close contact groups (Figure 1). In general, the lowest [IgG] were observed in the PREVAIL 3 contacts and the PREVAIL 1 placebo groups across all four assays (Figure 1A and C). As expected, highest [IgG] were detected in the PREVAIL 3 EVD survivor group across all four assays (Figure 1B). Some unusual values were notable: four EVD survivors had low [IgG] in at least one assay, and two of these had concordant results across all four assays (Figure 1B, leftmost values part); three placebo recipients displayed markedly higher [IgG], confirmed by all four assays (Figure 1C, rightmost values; Table S1.2).

**Figure 1.**
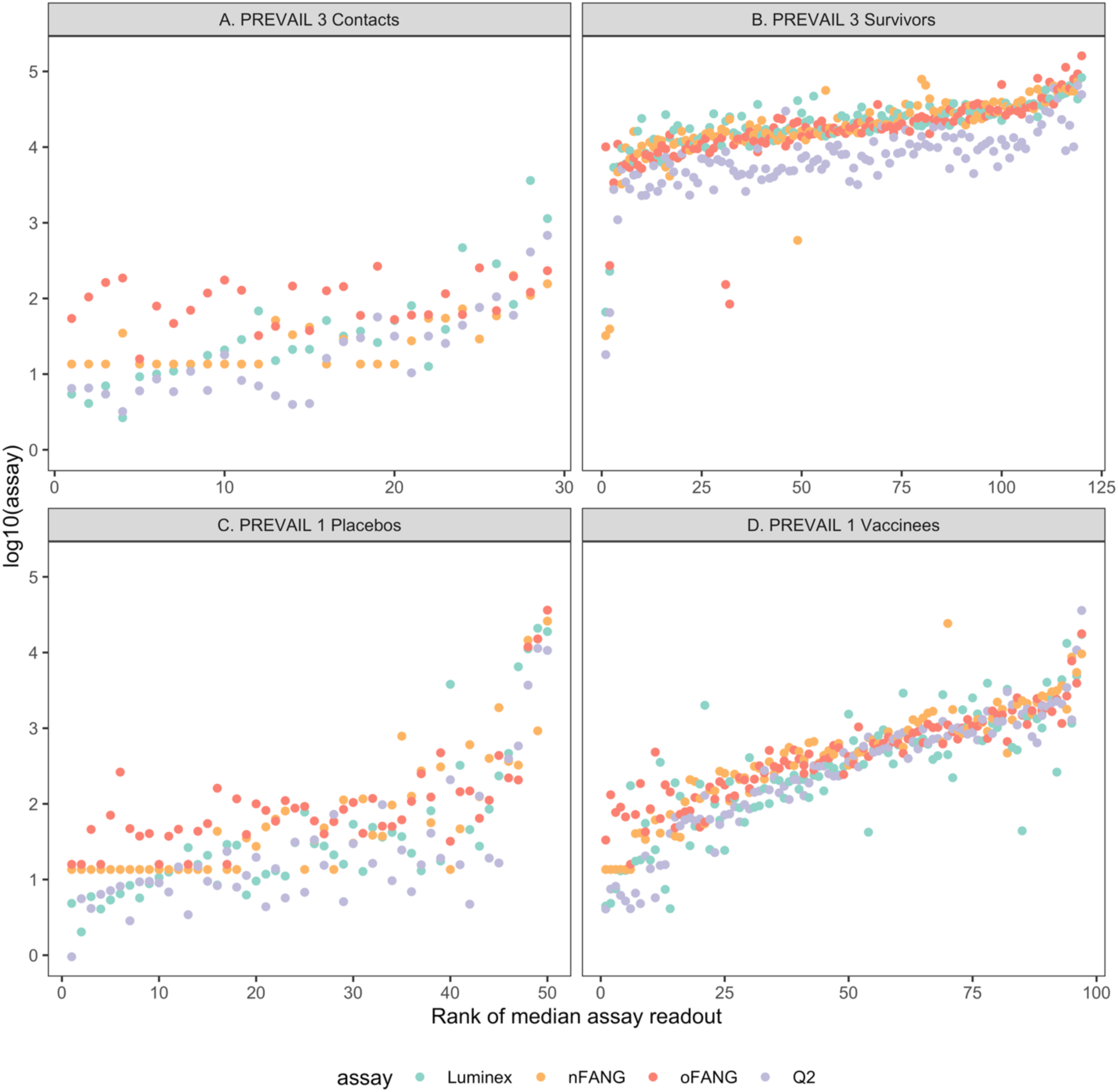
Individual assay readouts for the four anti-EBOV IgG assays in the close contact and survivor groups (PREVAIL 3), and placebo and vaccine groups (PREVAIL 1). Within each group, participants are ranked by the average of the four assays, and this rank is the x-axis value. Colors denote different assays.

### Assay linearity and conversion

Pairwise correlations among the four assays (Figure 2) showed that the log10 assay [IgG] readouts were linearly related and in reasonable agreement. The lowest Pearson correlation was 0.91 and was observed between the oFANG and the Luminex assays. The highest correlation was observed between the Q2 and the Luminex assays (0.95). In addition, slopes of the different pairs of assays can provide information about assay linearity. Commonly, to evaluate the linearity of an assay, serial dilutions of a sample are used. A regression of the log assay readout on the log of the dilution should have a slope of 1 over the linear range of an assay. It follows that, over the common linear range of two assays, a regression of paired readouts should also have a slope of 1. In our analyses, oFANG had slopes of 1.15 (95% CI: 1.12, 1.17), 1.25 (95% CI: 1.22, 1.27), and 1.15 (95% CI: 1.12, 1.17) for Q2, Luminex, and nFANG, respectively. The slopes for Luminex on Q2 and nFANG were both 1.08 (95% CI on Q2: 1.05, 1.12) (95% CI on nFANG: 1.05, 1.13), while the Q2 and nFANG assays had a slope of 0.99 (95% CI: 0.96, 1.02). Thus, the oFANG had slopes at least 10% different from the other assays, while the nFANG and Q2 slopes were within 1% of each other (Table 1). To visualize comparative performance, Figure S1.1 shows the Deming regression lines for the Luminex, Q2, and oFANG assay on the nFANG. Bland-Altman plots for the different pairs of assays are shown in Figure S1.2.

**Figure 2.**
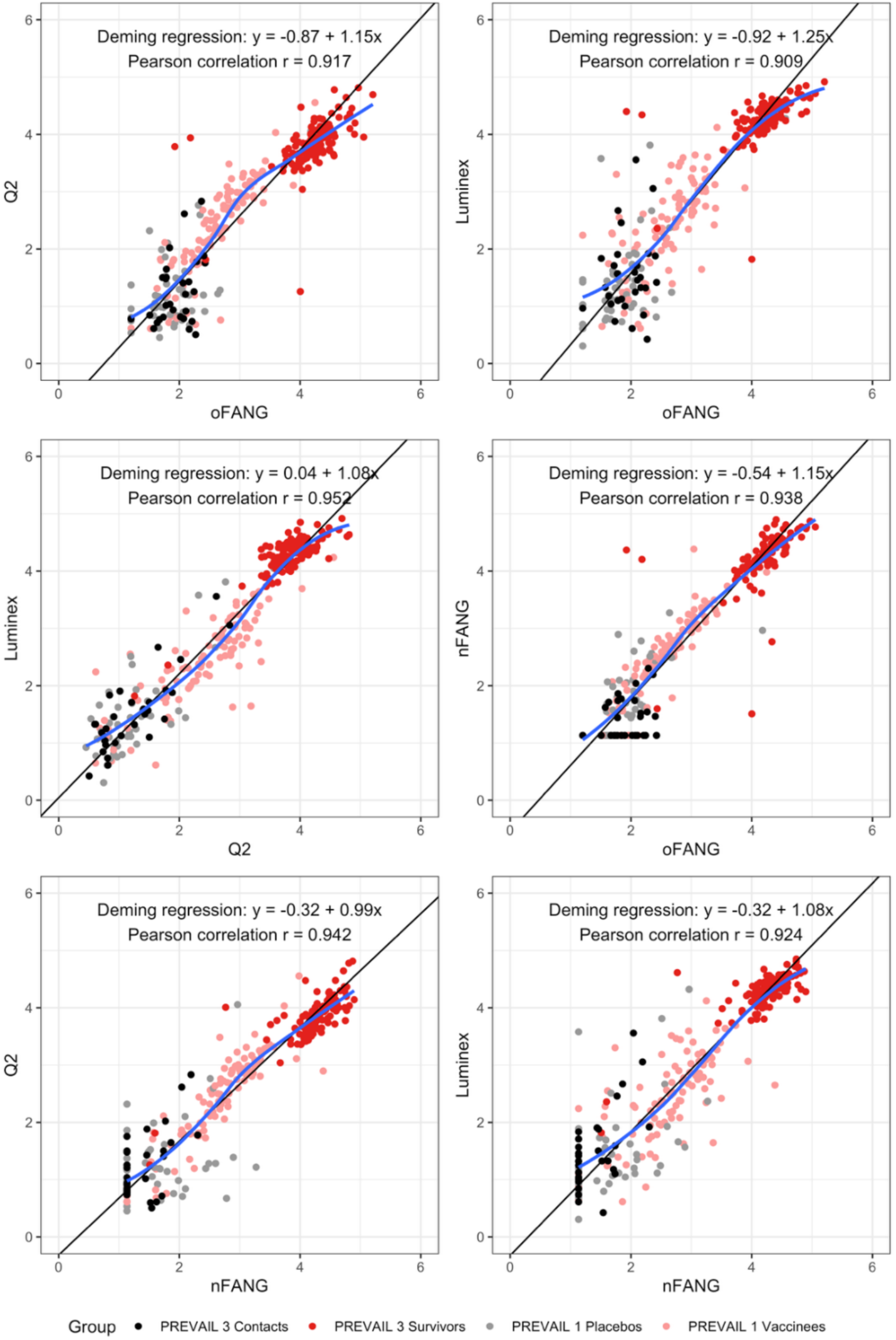
Pairwise correlations between the four anti-EBOV IgG assays. Colors indicate close contacts and survivors (PREVAIL 3), and placebos and vaccinees (PREVAIL 1).

**Table 1.**
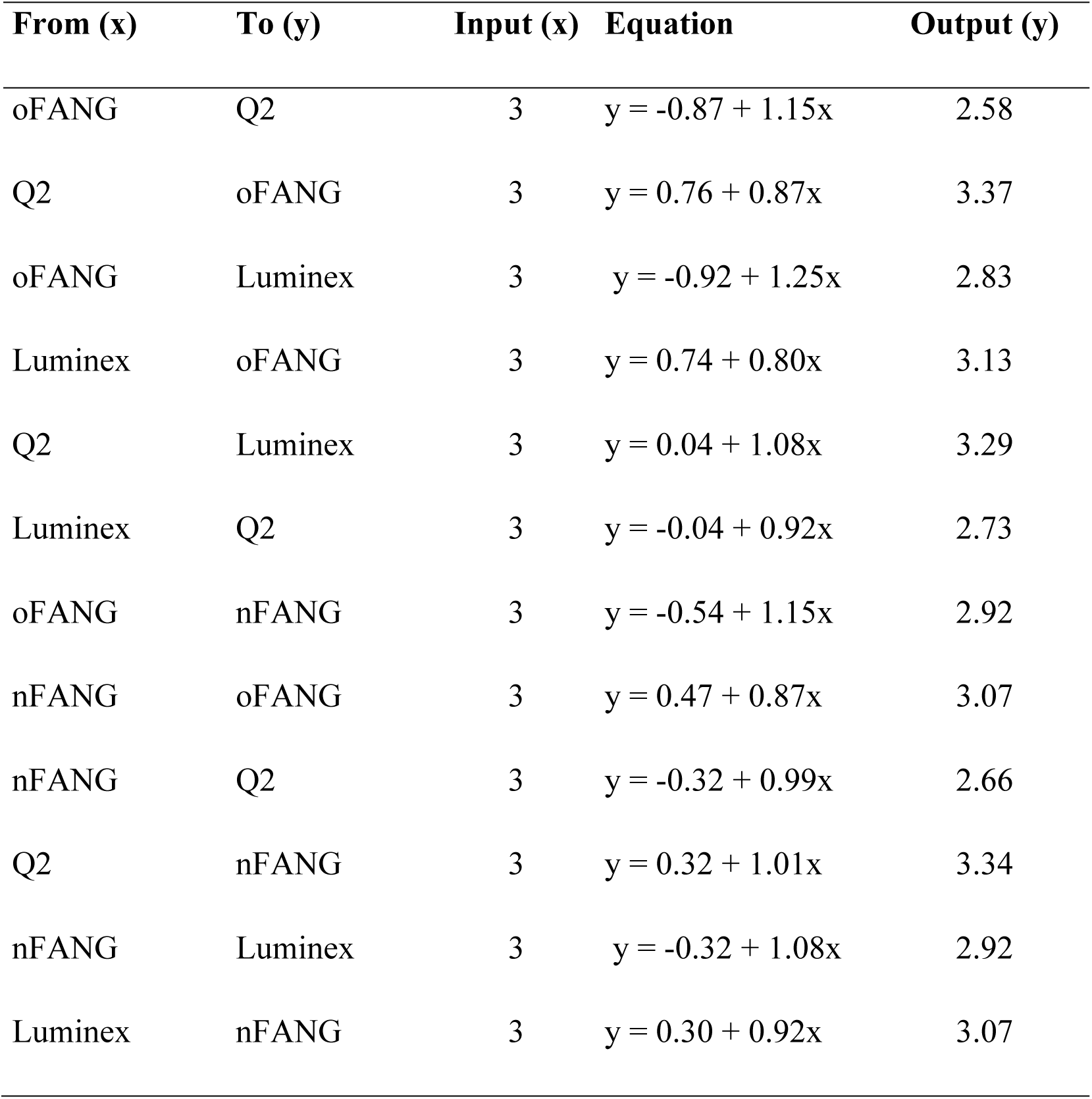
Conversion equations for four anti-EBOV IgG assays by Deming regression.

| From (x) | To (y) | Input (x) | Equation | Output (y) |
| --- | --- | --- | --- | --- |
| oFANG | Q2 | 3 | $y = -0.87 + 1.15x$ | 2.58 |
| Q2 | oFANG | 3 | $y = 0.76 + 0.87x$ | 3.37 |
| oFANG | Luminex | 3 | $y = -0.92 + 1.25x$ | 2.83 |
| Luminex | oFANG | 3 | $y = 0.74 + 0.80x$ | 3.13 |
| Q2 | Luminex | 3 | $y = 0.04 + 1.08x$ | 3.29 |
| Luminex | Q2 | 3 | $y = -0.04 + 0.92x$ | 2.73 |
| oFANG | nFANG | 3 | $y = -0.54 + 1.15x$ | 2.92 |
| nFANG | oFANG | 3 | $y = 0.47 + 0.87x$ | 3.07 |
| nFANG | Q2 | 3 | $y = -0.32 + 0.99x$ | 2.66 |
| Q2 | nFANG | 3 | $y = 0.32 + 1.01x$ | 3.34 |
| nFANG | Luminex | 3 | $y = -0.32 + 1.08x$ | 2.92 |
| Luminex | nFANG | 3 | $y = 0.30 + 0.92x$ | 3.07 |

Use of the Deming regression equations enabled conversion of [IgG] readouts across assays in this study (Table 1). For example, a readout of 3.00 EU/ml on the oFANG assay (Table 1 - Input) corresponds to 2.58 EU/ml on the Q2 assay (Table 1 - Output). Values below LOD were set to LOD/2. We also performed sensitivity analyses in which values below LOD were treated in two other ways: 1) we used the raw values; and 2) we removed paired values when either readout was less than the LOD. The parameter estimates were insensitive to how values below LOD were handled, and the conversion equations were robust (Table S1.3; Figure S1.3). In addition, re-run of the Luminex assay, removing 39 values that were above the ULOQ of the assay, was robust to the removal (Table S1.3).

### Performance of assays in distinguishing between two groups

When stratified by study (contacts versus EVD survivors for PREVAIL 3; vaccine versus placebo recipients for PREVAIL 1), all four assays discriminated well between the groups (Figure 3). The standardized mean difference, measuring the ability of an assay to discriminate between two groups, was similar across assays for the PREVAIL 1 groups (vaccine vs. placebo), with the Q2 assay showing the highest value (Table 2). For PREVAIL 3, the nFANG assay had the greatest ability to discriminate between EVD survivors and their close contacts, while Q2 and Luminex were notably lower (Table 2). In general, the standardized mean differences were much larger for the PREVAIL 3 groups across the assays, reflecting the higher EBOV GP_1,2_ [IgG] in EVD survivors compared to vaccine recipients. Greater variability was observed in PREVAIL 1 placebo recipients (standard deviations from 0.55 to 0.80) than the PREVAIL 3 close contacts (standard deviations from 0.37 to 0.47).

**Figure 3.**
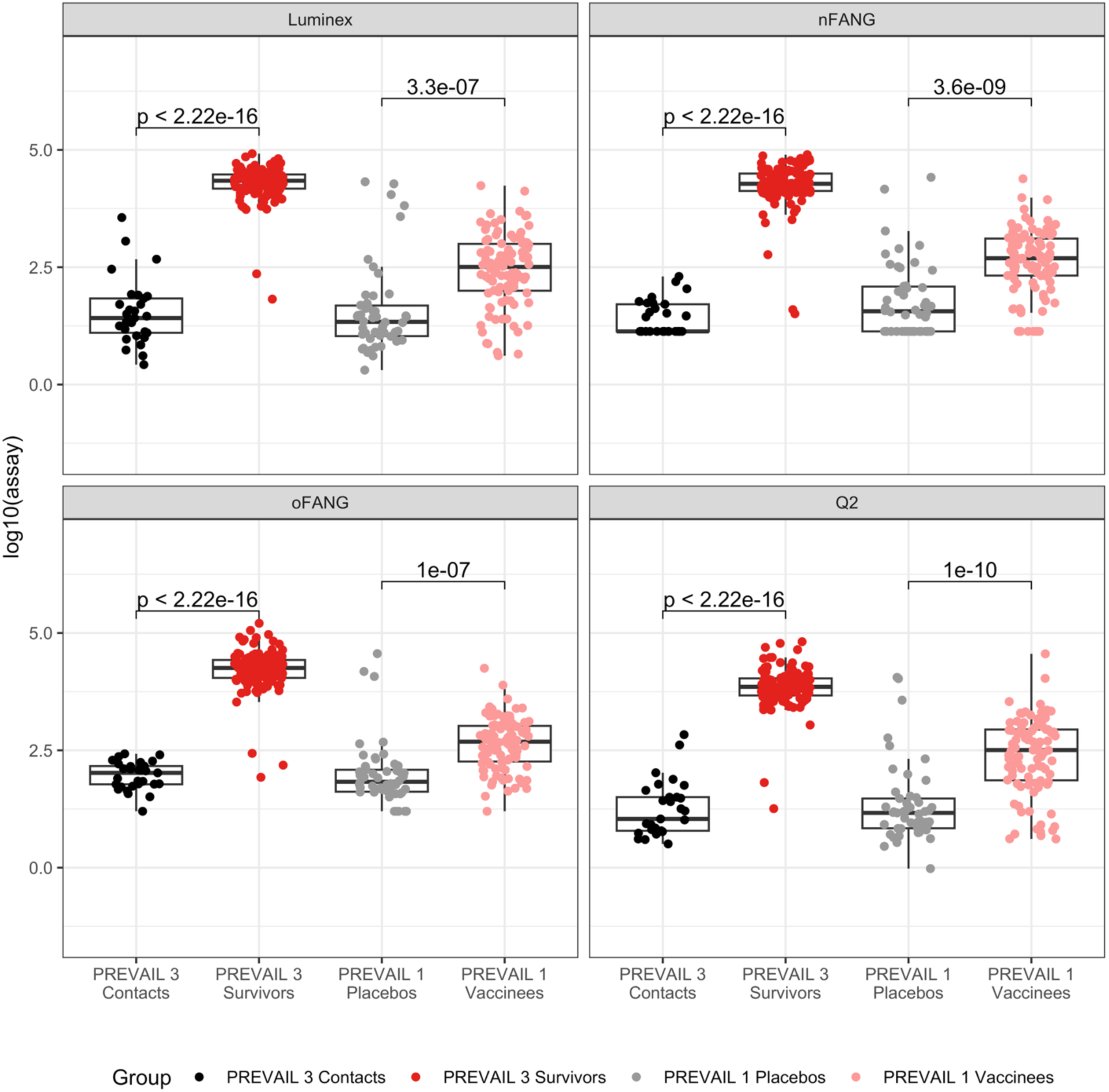
Box and whisker plots and t-test p-values s comparing contacts to survivors (PREVAIL 3) and placebos to vaccinees (PREVAIL 1) for the four anti-EBOV IgG assays.

**Table 2.** Mean of log10 assay values (standard deviations) and standardized mean differences (SMDs) (standard deviations) comparing placebo to vaccine recipients and survivors to contacts.

| **Assay** | **PREVAIL 1** |  |  | **PREVAIL 3** |  |  |
| --- | --- | --- | --- | --- | --- | --- |
|  | **Vaccine**<br>**(N=99)** | **Placebo**<br>**(N=50)** | **SMD** | **Survivors**<br>**(N=120)** | **Close**<br>**Contacts**<br>**(N=29)** | **SMD** |
| oFANG | 2.63 (0.54) | 1.98 (0.69) | 0.74 (0.21) | 4.22 (0.44) | 1.96 (0.30) | 4.23 (0.57) |
| nFANG | 2.64 (0.68) | 1.78 (0.79) | 0.83 (0.19) | 4.25 (0.47) | 1.43 (0.36) | 4.73 (0.77) |
| Q2 | 2.40 (0.80) | 1.34 (0.84) | 0.91 (0.20) | 3.84 (0.43) | 1.25 (0.59) | 3.54 (0.57) |
| Luminex | 2.44 (0.77) | 1.58 (0.94) | 0.70 (0.19) | 4.29 (0.37) | 1.53 (0.70) | 3.45 (0.68) |

To further quantify the ability of the assays to discriminate between groups, we estimated the ratio of sample sizes required to power a future study comparing two groups of participants for all four assays (Table 3), assuming the same repeat rate across assays. For a study comparing vaccine and placebo recipients, the sample size ratio’s confidence interval included one for all assay comparisons except for Luminex vs. Q2 for a vaccine study (67% larger sample size for Luminex), and for Q2 vs. nFANG for a survivor study (79% larger sample size for Q2).

**Table 3.**
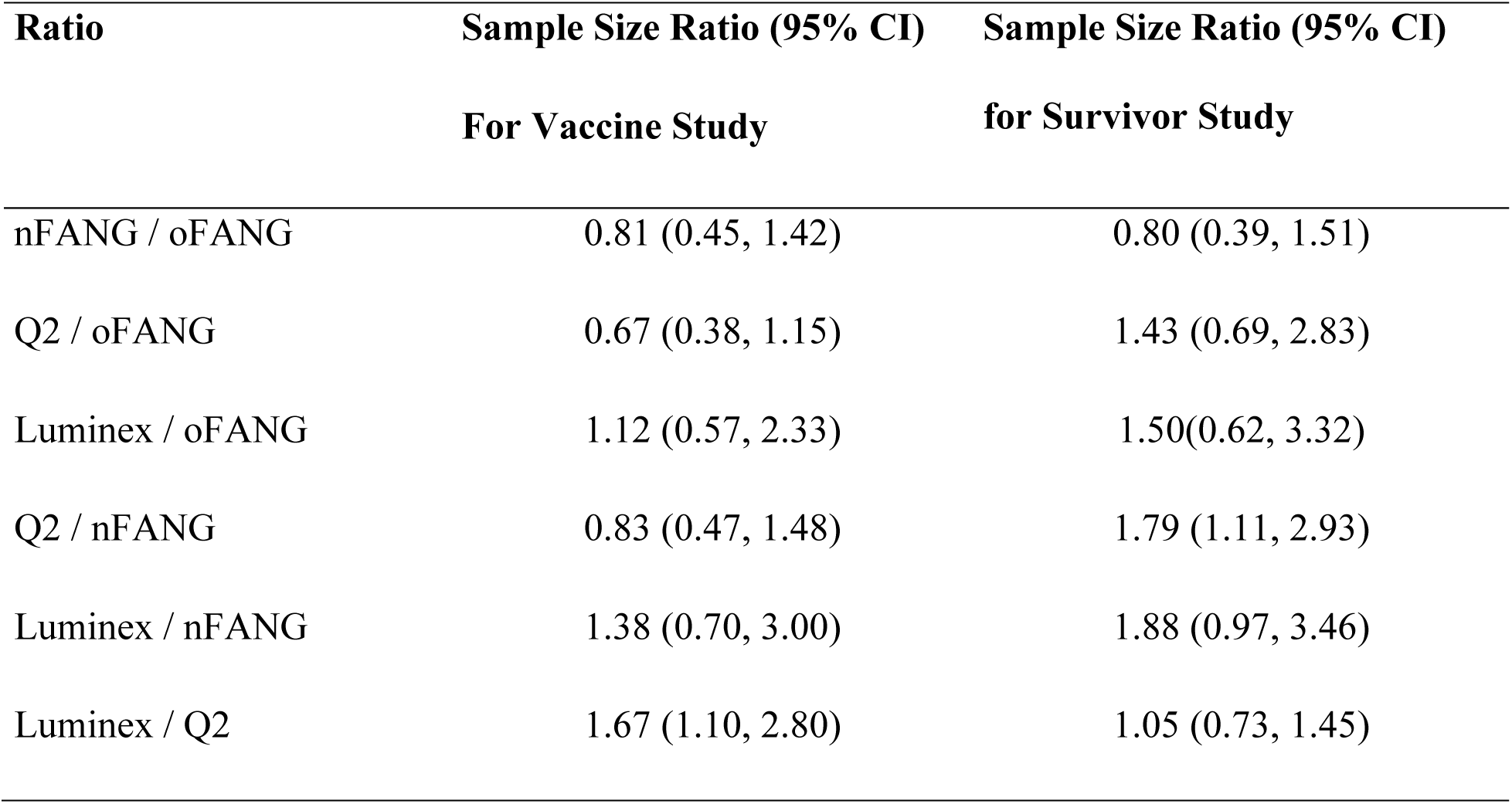
Sample size ratio (95% confidence interval) to power a study to detect a difference in mean EBOV GP_1,2_ antibody between groups. Sample size ratio for Assay 1/Assay 2 = 1.50 means Assay 1 requires 50% more participants.

| Ratio | Sample Size Ratio (95% CI) | Sample Size Ratio (95% CI) |
| --- | --- | --- |
|  | For Vaccine Study | for Survivor Study |
| nFANG / oFANG | 0.81 (0.45, 1.42) | 0.80 (0.39, 1.51) |
| Q2 / oFANG | 0.67 (0.38, 1.15) | 1.43 (0.69, 2.83) |
| Luminex / oFANG | 1.12 (0.57, 2.33) | 1.50(0.62, 3.32) |
| Q2 / nFANG | 0.83 (0.47, 1.48) | 1.79 (1.11, 2.93) |
| Luminex / nFANG | 1.38 (0.70, 3.00) | 1.88 (0.97, 3.46) |
| Luminex / Q2 | 1.67 (1.10, 2.80) | 1.05 (0.73, 1.45) |

### Other assay characteristics

Samples may need to be re-assayed due to calibrator or quality control (QC) failures, values falling below or above assay range, or replicate CV percentage values being above a pre-determined threshold (typically 20%). Sample failure is defined as failure to produce results even after repetition. The FANG assay has historically had high sample-repeat rates: an estimated 71.5% of sample measurements had to be repeated for a subset of data from the PREPARE study (23), with an estimated sample failure rate of 4%. This is much higher than the repeat rates (16.1% and 4%) and failure rates (0.7% and 2%) measured in our study for the Q2 and Luminex assays, respectively. However, it is important to note that Luminex repeat rates increased to 16.8% due to changes in the assay-passing algorithm in the second version of the assay, which was developed after our current analyses. In our study, repeat rates also differed between study type: for the Q2 assay, repeats due to an out-of-assay range were higher for PREVAIL 3 compared to PREVAIL 1, due to higher antibody titers in EVD survivors necessitating higher sample dilutions (Table S1.4).

In addition to higher sample repeat and failure rates, the FANG assays also had the lowest sample throughput rates, with only four to five samples per 96-well plate due to the required sample dilutions (Table 4). FANG throughput was further limited by the need to manually coat plates with the EBOV GP_1,2_ monomer as compared to the pre-printed Q2 plates. The Luminex assay required the least sample volume.

**Table 4.**
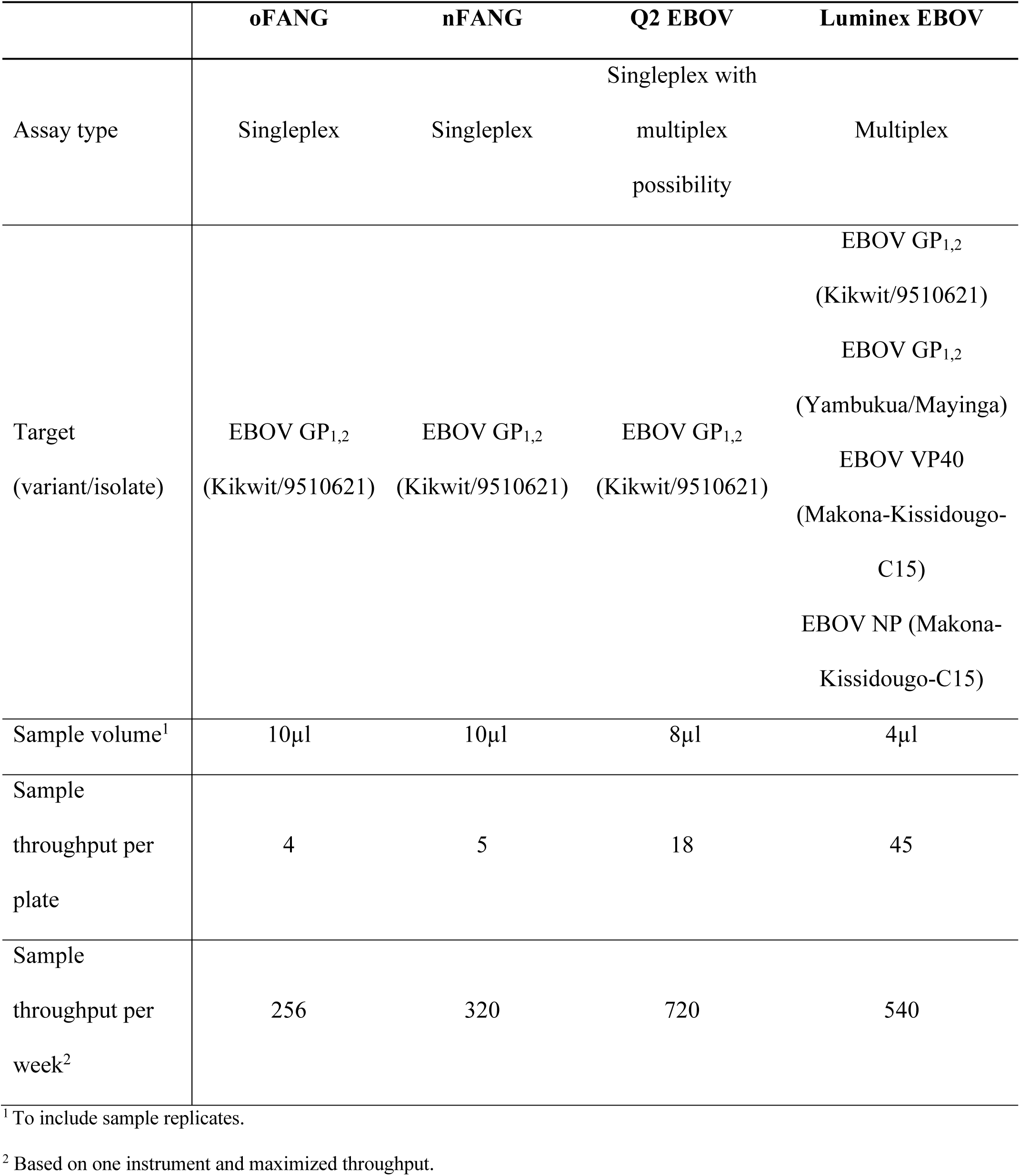
Comparison of the operating features of four anti-EBOV IgG assays.

The multiplex capability of the Luminex assay allowed for simultaneous antibody measurements for EBOV GP_1,2_ Yambuka/Mayinga, EBOV GP_1,2_ Kikwit/9510621, EBOV VP40, and EBOV NP, enabling a more comprehensive analysis of antibody responses to multiple viral antigens. Figures S1.4 and S1.5 show the performance of Luminex against all four target antibodies. The ability to distinguish between groups is lower with EBOV GP_1,2_ Yambuka/Mayinga compared to EBOV GP_1,2_ Kikwit/9510621, especially when comparing the placebo vs. vaccine arms (Figure S1.4, top panels). In addition, for EBOV VP40 or EBOV NP, there is discrimination between EVD survivors and their close contacts but no discrimination between vaccine and placebo recipients (Figure S1.4, lower panels), as expected. Figure S1.5 graphs the paired readouts for all six pairs of targets for the Luminex assay.

## DISCUSSION

EBOV glycoprotein is the sole external protein on the virion surface, and hence the primary target for antibody-based therapeutics and vaccines. EVD survivors typically develop robust anti-GP IgG responses, and the magnitude of these responses generally correlated with the level of protection against lethal infection in NHP vaccine studies (24, 25). Therefore, most ELISA-based assays measuring EBOV-specific antibody responses, whether induced by natural infection or vaccination, have targeted the detection of EBOV GP-specific IgG. As more advanced and higher-throughput assays are developed, the need to compare results across assays becomes more apparent. Whether for assay transition during ongoing long-term studies, or to inform decision making for upcoming studies, thorough validation, between-assay comparison, and within-study bridging analysis is critical to evaluating results.

The high correlation among the four assays assessed in our study (oFANG, nFANG, Q2, and Luminex) indicated similar performance for measuring anti-EBOV GP_1,2_-specific IgG responses across select groups. All four assays were concordant among the survivor, close contact, vaccine, and placebo groups, and all four distinguished EBOV antigen-exposed (survivors, vaccinated) and unexposed (contacts, placebos) groups with highly significant values. In addition, high between-assay correlations in our study allowed for the development of assay conversion equations, which may be useful for comparing assay readouts within a study (bridging).

However, the coefficients of these equations may vary depending on the laboratory, reagent batches, etc., and should ideally be estimated for each study. Evaluating assay interlaboratory variability was beyond the scope of this study.

Linear performance over some range of antibody concentration is an important assay feature (i.e., a sample that is diluted by 50% should have a readout that decreases by 50%). The slope of the regression of readouts on log dilution should be 1.0. If two assays are linear over the same range, the Deming slope should also be 1.0. For the assays we evaluated, the Deming slope for the Q2 and nFANG assays was virtually 1.0. The Luminex assay had a slope within 5% of 1.0 when compared to the nFANG and Q2 assays, while the oFANG had a slope greater than 10% of 1.0 for all other assays. These results suggest that the Q2 and nFANG assays have greater fidelity to linearity than the other assays; however, without performing a serial dilution experiment for all four assays, it is not possible to determine which assay(s) are truly closest to linear. Since the Luminex assay is only evaluated on a single sample dilution, these comparisons were not possible in our study.

Assay sensitivity and specificity are also critical. Because we did not have an *a priori* defined positivity threshold for the Q2 assay, we did not compare the assays in terms of this metric.

However, our estimates of the sample size required to power a future study indicated advantages of the Q2 assay over the Luminex assay for a placebo-controlled vaccine trial (67% larger sample size for Luminex), and for the nFANG over the Q2 assay for comparing EVD survivors vs. contacts (79% larger sample size for Q2).

Although our findings suggest that there is not a clear choice for the best assay in terms of sample size and power, pragmatic considerations, including assay cost, repeat rate, and algorithm complexity, also play a role in assay selection. One clear benefit of the Luminex assay is the ability to multiplex, simultaneously measuring not only anti-EBOV GP_1,2_ antibodies against multiple EBOV isolates, but also antibodies elicited against EBOV NP and VP40. Including multiple EBOV isolates of GP_1,2_ may provide insights into cross-reactivity and cross-protection. Versus the FANG assays, another benefit of the Luminex and Q2 assays is their increased throughput. In contrast to the multiple sample dilutions required by the FANG assays, minimal dilutions required by Luminex and Q2 assays allow more samples to be run on the same plate. The overall lower repeat rate of these two assays compared to FANG assays also increases throughput and likely decreases the overall cost of assay execution. However, as we show, sample repeat rates may vary based on study type and the proportion of samples from participants with high-antibody titers.

During the 2013–2016 Western African EVD outbreak, mainly due to limited access to specialized equipment, laboratory infrastructure, and expertise, some ELISA-based anti-EBOV GP assays were performed in laboratories outside of Africa. Since then, laboratory capability in Western and Central Africa has developed significantly, with several laboratories in EVD at-risk countries now able to perform ELISA assays (12, 26). Luminex-based assays have also been transferred to EVD outbreak regions (27). Nonetheless, challenges still exist in these often resource-limited settings, and, in addition to assay cost, assay and equipment complexity may also have to be taken into consideration. Bead-oriented Luminex platforms with complex optical and fluidics systems require meticulous and precise calibration and equipment maintenance, which may be more difficult to achieve in hard-to-reach areas.

In summary, we find that all four compared assays are suitable for detecting EBOV anti- GP_1,2_ IgG antibody responses. Therefore, assay choice for future studies may be determined by assay throughput, cost, variability, and complexity, particularly where sample availability is limited and in more austere settings.

## METHODS

### Study design

Samples for assay comparison were selected from two clinical studies conducted in Liberia in response to the 2013–2016 Western African EVD outbreak caused by the EBOV Makona variant (28). PREVAIL 1 (NCT02344407) was a randomized, placebo-controlled, double-blinded Phase 2 clinical trial of two EBOV vaccines, rVSVΔG-ZEBOV-GP (Kikwit/9510621 EBOV variant/isolate) and ChAd3-EBO-Z (Yambukua/Mayinga EBOV variant/isolate), involving 1,500 participants without a history of EVD (9). PREVAIL 3 (NCT02431923) was a longitudinal observational cohort study of 3,930 Liberian EVD survivors and their close contacts for five years (10).

From PREVAIL 1, 149 samples were selected from month 36 post-vaccination until the end of longitudinal follow-up. Samples were randomly selected with stratification by vaccination group and study visit. From PREVAIL 3, 149 samples were selected, including half from baseline visits and half from the one-year follow-up visits. At each visit, 20% of the samples from PREVAIL 3 were from seronegative contacts, with the remainder from seropositive EVD survivors. Within each of the above groups of PREVAIL 3 participants, specimens were randomly selected and stratified by the distribution quartiles of known oFANG results.

### Assays and data

A total of 298 frozen serum samples from the PREVAIL 1 and PREVAIL 3 studies were transported from the Liberia Institute for Biomedical Research in Monrovia, Liberia, to the Integrated Research Facility at Fort Detrick (IRF-Frederick), National Institute of Allergy and Infectious Diseases (NIAID), in Frederick, MD, USA. An aliquot of these samples was then shipped to the French National Research Institute for Sustainable Development (IRD) laboratory, Montpellier, France. The nFANG and Q2 assay testing was conducted at IRF-Frederick.

Luminex Version 1 assay testing was conducted at the French National Research IRD. The validation reports for the Q2 and the Luminex Version 1 assays are provided in the Supplemental Materials. Each assay was calibrated using its own platform-specific standards: the Luminex assay used the WHO International Standard 15/220, and the FANG and Q2 assays used an internal reference standard (BMIZAIRE108) previously correlated to the WHO 15/220 (7).

### oFANG

Anti-EBOV GP_1,2_ IgG titers previously measured on the 298 samples using the oFANG ELISA, following the qualified FANG protocol, were obtained from the University of Minnesota PREVAIL database (9, 10). Briefly, Immulon 2 HB 96-well microplates (Thermo Fisher Scientific, Walkersville, MD, USA) were coated with recombinant EBOV GP_1,2_. Test samples (replicates), quality controls (6-point, 1:2 dilution series), the reference standard (11-point, 1:2 dilution series), and a negative control (1:50 dilution, commercially obtained pooled naïve human serum from U.S. donors, Sera Care, Milford, Ma) were added to the plates, followed by incubation, washes, and the addition of the horseradish peroxidase-conjugated, anti-human IgG (Thermo Fisher Scientific, Walkersville, MD, USA), the TMB substrate (Thermo Fisher Scientific, Walkersville, MD, USA), and the stop solution. Plates were read on a SpectraMax Plus 384 plate reader (Molecular Devices, Sunnyvale, CA, USA) at 450 nm. Well background optical densities were measured at 650 nm and subtracted from the 450-nm optical density readings.

The oFANG assay readout was obtained as follows: If the sample CV among dilutions was > 20%, the dilution furthest from the mean was masked until the CV fell below 20% and the log10 average was used. If the CV remained above 20% or too many dilution points were masked, the sample was repeated. If fewer than three unmasked dilutions met CV criteria or if the inter-replicate CV exceeded 30%, the sample was repeated once. If no acceptable values were obtained, sample failure was assumed, and the result was recorded as missing data.

### nFANG

Anti-EBOV GP_1,2_ IgG titers on all 298 samples were measured using the nFANG ELISA, following the validated FANG protocol, as previously described (7, 12) (detailed in the Supplemental Materials). Briefly, Immulon 2 HB 96-well microplates (Thermo Fisher Scientific, X1530419 NT521019) were coated with the recombinant EBOV GP_1,2_ and then loaded with the reference standard (BMIZAIRE108, 11-point, 1:2 dilution series), samples (1:50, 1:100, 1:200, 1:400, 1:800, and 1:1,600 dilutions; sample replicates were ran on separate plates), quality controls (duplicates, 8-point dilution series), and a negative control (commercially obtained pooled naïve human serum from U.S. donors, Sera Care, Milford, MA, USA). After adding horseradish peroxidase-conjugated anti-human IgG secondary antibody (Thermo Fisher Scientific, Walkersville, MD, USA), the TMB substrate (Thermo Fisher Scientific, Walkersville, MD, USA), and the stop solution, with incubations and washing, the plates were read using the SpectraMax Plus 384 colorimeter plate reader (Molecular Devices, Sunnyvale, CA, USA) at 450 nm; well background optical densities at 650 nm were subtracted using Softmax 7.1 (Molecular Devices, Sunnyvale, CA, USA).

The nFANG assay readout was obtained as follows: If the CV between replicates on each plate was ≤ 30%, the log10 average was used; otherwise, that dilution was masked. If the CV between unmasked dilutions was ≤ 20%, the log10 average was used; otherwise, dilutions were masked until an acceptable CV was achieved. If fewer than three unmasked dilutions met CV criteria, or if the inter-plate CV exceeded 30%, the assay was repeated once. If all values were above the assay range, higher dilutions were tested (1:2,000, 1:4,000, 1:8,000, 1:16,000, 1:32,000, and 1:64,000); additional dilutions (1:250, 1:500 and 1:1,000) were run per need. If no dilutions met CV criteria after one repeat, sample failure was assumed and recorded as missing data.

### Q2

Anti-EBOV GP_1,2_ IgG titers were also measured on the same 298 samples using the EBOV GP SIMOA Planar assay on an SP-X Automated Immunoassay Analyzer (Quanterix) (detailed in the Supplemental Materials). Briefly, for the Q2 assay, calibrator standard (BMIZAIRE108, seven-point 1:3 serial dilution), high-concentration and low concentration quality controls (1:4,000 dilution), and study samples (serial dilutions of 1:50 and 1:4,000) were added in duplicates to 96-well plates pre-coated with recombinant EBOV GP. Following washes and incubations, and the addition of the Streptavidin-HRP antibody and Super Signal® Substrate, the plates were imaged on the SP-X and data was processed via the SP-X analysis application.

The Q2 assay readout was obtained as follows: If both sample dilutions had a replicate CV ≤ 20%, and dilutions were within 30% of each other, the log10 of the average was used. If only one dilution had a passing CV, that value’s log10 was used. If no dilutions passed, the assay was repeated once. If values for both dilutions were above the assay range, additional dilutions (1:6,000 or 1:8,000) of the sample were tested. If acceptable CV criteria were met at higher dilutions, the log10 average or log10 of a single passing dilution was used. If no dilutions were satisfactory after one repeat, sample failure was assumed and counted as missing data. If the 1:50 dilution fell below the LOD (0.0009 EU/mL), the value was imputed as LOD/2.

### Luminex EBOV Kikwit GP Assay

Anti-EBOV GP_1,2_ IgG titers were measured in the aliquots of the 298 samples using a Luminex-based Version 1 protocol described previously (16) that was updated with GP_1,2_ Kikwit/9510621, which is used in certain vaccines (Supplemental Materials). Briefly, recombinant EBOV proteins (NP, VP40, GP_1,2_ Kikwit/9510621 and GP_1,2_ Yambukua/Mayinga) were coupled to vortexed carboxy-functionalized beads, diluted to 2,000 beads/μl, and added to the 96-well flat-bottom chimney plates (Greiner bio one, Frickenhausen, Germany). Following aspiration using an automatic plate washer (BioTek 405TS Microplate washer), the reference standard (WHO 15/220), quality controls, and samples (1:200 dilution, duplicates) were added. After incubation and washes, and the addition of biotin-labeled anti-human IgG (BD-Pharmingen, Le Pont De Claix, France) and streptavidin-R-phycoerythrin (Fisher Scientific/Life Technologies, Illkirch, France), the antigen-antibody reactions were read on Intelliflex (Luminex Corp., Austin, TX, USA). At least 100 events were read per bead set, and the results were expressed as median fluorescence intensity (MFI) per 100 beads.

The Luminex assay readout was obtained as follows: If Net MFI values for duplicates had a CV < 20%, the log10 of the average was used. If the CV exceeded 20% and the average MFI was > 300, the results were invalid, and the sample was retested in duplicate. If both rounds failed due to high CV, the sample was considered invalid and counted as missing data. Values exceeding the ULOQ were not repeated and were analyzed as is. In the updated Version 2 of the assay, developed after this study, samples above the assay range were repeated at appropriate dilutions.

### Statistical Methods

Antibody concentrations were log10 transformed and summarized using means and standard deviations. Because the Luminex assay used a different reference standard (WHO 15/220) compared to the FANG and Q2 assays (BMIZAIRE108), all Luminex values were converted to EU/ml according to a predetermined correlation (Luminex (EU/mL) = GP_1,2_ Kikwit/9510621 (IU/mL)*200*27135.9) (7). Deming regressions and Pearson correlations were used to estimate the relationships between all pairs of the four assays. The Deming regression equations were used to convert the reading of one assay to an equivalent value in another assay. Values that were below the LOD were set to LOD/2. Sensitivity analyses removed values below LOD or used the actual readouts (even if below LOD). A listing of the LOD and LLOQ values used in the calculations is provided in the Supplemental Materials (Table S1.5).

T-tests were used to compare means between groups (placebo vs. vaccine recipients in PREVAIL 1; EVD survivors vs. contacts in PREVAIL 3). To quantify the overall ability of an assay to discriminate between two groups, we calculated the standardized mean difference (SMD), i.e., the difference in sample means divided by the square root of the sum of the sample variances for the two groups. The square of the SMD is proportional to the sample size needed to power a future study with equal allocation to each group and thus a measure of its ability to distinguish between two groups. Thus, if the squared ratio of the SMDs for two assays is 1.50, then 50% more participants are needed for the assay with the smaller SMD to achieve a given power. Sample-size ratios were estimated for all possible pairs of assays in both PREVAIL 1 and PREVAIL 3 comparisons. Confidence intervals for the sample-size ratios were calculated using the nonparametric bootstrap via the percentile method with 10,000 bootstrap samples. All tests were two-sided, with a type I error rate of 0.05 considered significant. There was no correction for multiple comparisons. All data analyses were conducted in R Statistical Software version 4.3.0.

### Study approval

All of the PREVAIL 1 and PREVAIL 3 samples were collected and stored according to the IRB-approved language in each protocol governing their future use in Ebola-related research. The samples were collected only after written informed consent was obtained from each participant. Subsequent to completion of the antibody titer assay development work described in this manuscript, the NIH IRB determined that the specific nature of the work should have fallen under the FDA policy for in vitro diagnostic (IVD) assay development and should either have been filed under an IVD with FDA or received a prospective IRB-approved exemption to the policy. In making this determination, the IRB declared therefore that the work was noncompliant with their oversight, but that this infraction was neither serious nor continuing, and that no further action was needed. The manuscript was subsequently approved for submission and publication.

## Data availability

Data can be made available from the corresponding authors as it is human-subject derived. Values for all data points in graphs are reported in the Supporting Data Values file.

## AUTHOR CONTRIBUTIONS

D.F., C.R., I.M.B., C.R., N.D.C., Y.L, and H.C.L conceptualized the study, P.C., A.J., H.G., S.C., N.B., A.A., M.P.(Peeters Martine), G.T.(Guillaume Thaurignac), N.Bi.(Neraide Biai) supervised and conducted laboratory experiments, S.B.F., V.C., E.L. performed statistical analyses, D.F., C.R., I.M.B., C.R., N.D.C., E.L., P.C, N.B., and L.R. interpreted study results. All authors contributed to writing and editing the manuscript.

## FUNDING SUPPORT

This work was supported by NIAID, NIH. This work was supported in part through a Laulima Government Solutions, LLC, prime contract with the NIH NIAID (HHSN272201800013C). I.M.B., P.C., A.J., and H.G. performed this work as employees of Laulima Government Solutions, LLC. This project has been funded in whole or in part with federal funds from the National Cancer Institute, National Institutes of Health, under Contract No. 75N91019D00024. PREVAC-UP is part of the EDCTP2 programme supported by the European Union (grant number RIA2017S-2014 – PREVAC-UP).

## ACKNOWLEDGMENTS

We thank Anya Crane (NIH NIAID IRF-Frederick) and Rachel Nowak for critically editing the manuscript. The views and conclusions contained in this document are those of the authors and should not be interpreted as necessarily representing the official policies, either expressed or implied, of the U.S. Department of Health and Human Services or of the institutions and companies affiliated with the authors, nor does mention of trade names, commercial products, or organizations imply endorsement by the U.S. Government.

## Notes

### Competing Interest Statement

The authors have declared no competing interest.

