## Supplemental Material for "Comparison of Four Assays That Measure Antibodies to Ebola Virus Glycoprotein"

#### 1. ASSAY COMPARISONS

##### MATERIALS AND METHODS

**nFANG**

Anti-EBOV $GP_{1,2}$ IgG titers on all 298 samples were measured at IRF-Frederick using the FANG ELISA following the NFANG protocol, as previously described (1) (2). In detail, microtiter plates (Thermo Fisher, X1530419 NT521019) were coated with the recombinant Ebola glycoprotein and allowed to incubate at 4 °C for at least 14 hours in the absence of light. The starting dilution of 1:62.5 was prepared using an ELISA Diluent (ED) containing 5% weight-by-volume dry milk (LabScientific, Highlands, NJ, USA) and 0.1% volume-by-volume Tween-20 (FANG wash buffer) (Sigma Aldrich, St. Louis, MO, USA) in phosphate buffered saline (Gibco, Gaithersburg, MD, USA). The ED was used to prepare the reference standard (calibrator) in a dilution block as an 11-point, 1:2 dilution series. In a different dilution box, 5 µL of human serum samples were diluted in 495 µL of ED in duplicates, and an 8-point dilution series was performed. The same dilution series was used for high and low concentration quality controls (Battelle), and pooled naïve human serum (Sera Care, Milford, Ma) was used as a negative control. Thawed EBOV GP-coated plates were washed three times with 300 µL phosphate buffered saline with 0.1% volume-by-volume Tween-20, and 100 µL per well of the serially diluted reference standard, samples (sample replicates plated on different plates), and controls were added and incubated for one hour in 37 °C with absence of light. The plates were subsequently washed three times with FANG wash buffer followed by addition of 100 µL per well of the horseradish peroxidase-conjugated anti-human IgG secondary antibody (Thermo Fisher Scientific, Walkersville, MD, USA). Plates were incubated for another hour at 37 °C in absence of light before washing again three times with 300 µL of FANG wash buffer and adding TMB substrate (Thermo Fisher Scientific, Walkersville, MD, USA). Following a 30-minute incubation at room temperature in the absence of light, 100 µL of stop solution - sulphuric acid (Thermo Fisher Scientific, Walkersville, MD, USA) was added. Plates were measured in a SpectraMax Plus 384 colorimeter plate reader (Molecular Devices, Sunnyvale, CA, USA) at 450 nm. Individual well background optical densities were measured at 650 nm and subtracted from the 450 nm optical density using the Softmax 7.1 version software (Molecular Devices, Sunnyvale, CA, USA).

**Q2**

In addition, anti-EBOV $GP_{1,2}$ IgG titers were measured on the same 298 samples using the newly developed Ebola GP Simoa Planar assay on an SP-X Automated Immunoassay Analyzer (Quanterix). For the Simoa assay, kits were removed from the refrigerator and brought to room temperature at least 30 minutes before utilization. The 1X wash buffer provided in the kit was used to wash the 96-well plates pre-coated with recombinant Ebola $GP_{1,2}$ (at a titer of 400 µg/mL) on the Simoa™ Microplate Washer. A calibrator standard (provided in the kit) at a seven-point 1:3 serial dilution was created, and high concentration and low concentration quality controls (QCs - provided in the kit) were diluted with sample diluent (provided in the kit) at a 1:4000. Study samples were diluted with the kit-provided sample diluent and measured at serial dilutions of 1:50 and 1:4000. The calibrator, QCs, and samples were added to the plates in duplicates followed by an initial two-hour incubation with a hydrolyzed microcline lid on the Simoa™ Microplate Shaker at 23 °C and an orbital rotation of 525 RPMs. After incubation, the plates were washed, and a biotinylated antibody was added followed by a 30-minute incubation on the Simoa™ Microplate Shaker as described above. After a second plate wash, the Streptavidin-HRP antibody was added to the plates followed by a 30-minute incubation on the Simoa™ Microplate Shaker and a final plate wash. Super Signal® Substrate was made using 3mL of super signal substrate and 3mL Luminol Enhancer , and 50 µL of Super Signal® Substrate was added to each well. The glass bottom of each plate was wiped with a lint-free cloth and the plates were read within four minutes of adding the Super Signal® Substrate using the imager. Captured images were transferred to a USB drive and uploaded to the SP-X Analysis application.

**Luminex Kitwik GP Assay**

An aliquot of all 298 samples was shipped to IRD Montpellier TransVIHMI (France) to detect anti-EBOV $GP_{1,2}$ IgG antibodies using a Luminex-based protocol described previously (Ayouba et al, JCM, 2017) that was updated with an additional $GP_{1,2}$ antigen, GPkikwit95, used in certain vaccines. To that end, we coupled recombinant EBOV proteins (NP, VP40, GP kikwit95 and GP Mayinga 76) to carboxy-functionalized beads at a ratio of 2µg of recombinant proteins per million of beads. Before use, recombinant protein-coupled beads were vortexed for 30 seconds and diluted to 2,000 beads/µl of assay buffer (phosphate buffered saline (PBS)) containing 0.75 mol/L NaCl, 1% (wt/vol) bovine serum albumin (Sigma Aldrich, Saint-Quentin Fallavier, France), 5% (vol/vol) heat-inactivated fetal bovine serum (Gibco-Invitrogen, Cergy Pontoise, France), and 0.2% (vol/vol) Tween-20 (Sigma-Aldrich)). Tests were performed in 96-well flat-bottom chimney plates (Greiner bio one, Frickenhausen, Germany) and 50 µl of bead mixture was added to each well. The supernatant was aspirated using an automatic plate washer (BioTek 405TS Microplate washer). Next, the wells were incubated with 100 µL of plasma (diluted 1/200 in assay buffer) for 16 hours at 4˚C in the dark on a plate shaker at 400 rpm/min. After five washings with 100 µL of assay buffer, 50 µLl of biotin-labeled anti-human IgG was added (BD-Pharmingen, Le Pont De Claix, France) at a concentration of 4 µg/mL in each well and incubated for 30 minutes in the dark with continuous shaking at 400 rpm/min. Plates were washed five times as above, and 50 µL of streptavidin-R-phycoerythrin (Fisher Scientific/Life Technologies, Illkirch, France) at 4 µg/ml was added per well and incubated for 10 minutes while shaking at 400 rpm/min. After an additional five washes, 150 µL of reading buffer (PBS 1X, 1% de BSA) was added per well to read antigen-antibody reactions on Intelliflex (Luminex Corp., Austin, TX, USA). At least 100 events were read for each bead set, and the results were expressed as median fluorescence intensity (MFI) per 100 beads.

**Limits of Detection and Quantitation**

Each assay has its specific limit of detection (LOD), lower limit of quantitation (LLOQ), and upper limit of quantitation (ULOQ), but they are not comparable due to different units and the impact of these limits in any sample analyses is very much dependent on the handling of dilutions. We provide them below for accessibility (Table S1.5) but comparing limits across assays is problematic.

**TABLES AND FIGURES**

Table S1.1. Participant demographics.

| Characteristic | PREVAIL 3 contacts N = 29[1] | PREVAIL 3 survivors N = 120[1] | PREVAIL 1 placebo arm N = 50[1] | PREVAIL 1 vaccine arm N = 99[1] |
|---|---|---|---|---|
| Female, n (%) | 17 / 29 (59%) | 66 / 120 (55%) | 15 / 50 (30%) | 28 / 99 (28%) |
| Age, mean (SD) | 28 (15) | 34 (14) | 34 (13) | 34 (11) |

Table S1.2. Three participants with high values in all four assays. All participants were from the placebo arm of PREVAIL 1.

| Female | Age | nFANG GP | oFANG GP | Q2 GP | Luminex GP Kikwit/ 9510621 | Luminex GP Yambukua/ Mayinga | Luminex NP | Luminex VP40 |
|---|---|---|---|---|---|---|---|---|
| 0 | 22 | 4.163 | 4.073 | 3.568 | 3.296 | 3.416 | 3.961 | 3.882 |
| 0 | 19 | 2.965 | 4.180 | 4.055 | 3.522 | 3.687 | 3.957 | 3.365 |
| 1 | 21 | 4.416 | 4.560 | 4.028 | 3.486 | 3.554 | 3.799 | 4.004 |

Table S1.3. Deming regression estimates of slope and intercept using different methods for handling readouts that are below the limit of detection. 95% CIs are from 1000 bootstrap samples.

| y | x | method | intercept | Intercept 95%CI | slope | Slope 95%CI |
|---|---|---|---|---|---|---|
| Q2 | oFANG | LOD/2 | -0.871 | (-0.96, -0.77) | 1.150 | (1.12, 1.17) |
| Q2 | oFANG | raw values | -0.980 | (-1.00, -0.81) | 1.181 | (1.13, 1.18) |
| Q2 | oFANG | removing below LOD | -0.955 | (-0.99, -0.76) | 1.174 | (1.12, 1.18) |
| Luminex | oFANG | LOD/2 | -0.920 | (-1.07, -0.76) | 1.251 | (1.22, 1.27) |
| Luminex | oFANG | raw values | -0.972 | (-1.11, -0.81) | 1.267 | (1.23, 1.29) |
| Luminex | oFANG | removing below LOD | -1.007 | (-1.14, -0.82) | 1.275 | (1.23, 1.30) |
| Luminex | oFANG | removing above ULOQ | -0.983 | (-1.16, -0.84) | 1.280 | (1.24, 1.32) |
| Luminex | Q2 | LOD/2 | 0.042 | (-0.12, 0.12) | 1.083 | (1.05, 1.12) |
| Luminex | Q2 | raw values | 0.098 | (-0.12, 0.12) | 1.066 | (1.05, 1.12) |
| Luminex | Q2 | removing below LOD | 0.098 | (-0.12, 0.12) | 1.066 | (1.05, 1.12) |
| Luminex | Q2 | removing above ULOQ | 0.125 | (-0.10, 0.15) | 1.047 | (1.03, 1.12) |
| nFANG | oFANG | LOD/2 | -0.544 | (-0.60, -0.40) | 1.154 | (1.12, 1.17) |
| nFANG | oFANG | raw values | -0.530 | (-0.54, -0.36) | 1.153 | (1.11, 1.16) |
| nFANG | oFANG | removing below LOD | -0.158 | (-0.30, -0.10) | 1.057 | (1.03, 1.09) |
| Q2 | nFANG | LOD/2 | -0.316 | (-0.41, -0.17) | 0.993 | (0.96, 1.02) |
| Q2 | nFANG | raw values | -0.385 | (-0.48, -0.23) | 1.010 | (0.98, 1.04) |
| Q2 | nFANG | removing below LOD | -0.557 | (-0.61, -0.30) | 1.053 | (0.99, 1.07) |
| Luminex | nFANG | LOD/2 | -0.323 | (-0.53, -0.18) | 1.083 | (1.05, 1.13) |
| Luminex | nFANG | raw values | -0.344 | (-0.61, -0.25) | 1.085 | (1.06, 1.15) |
| Luminex | nFANG | removing below LOD | -0.647 | (-0.93, -0.42) | 1.159 | (1.11, 1.23) |
| Luminex | nFANG | removing above ULOQ | -0.288 | (-0.49, -0.20) | 1.068 | (1.05, 1.11) |

Table S1.4. Q2 assay repeat and sample failure rates per study.

| Type | PREVAIL 1 | PREVAIL 3 |
|---|---|---|
| Repeat rate: Out of assay range | 0.0% | 3.4% |
| Repeat rate: CV | 6.0% | 3.4% |
| Repeat rate: QC | 10.0% | 9.4% |
| Repeat rate: Total | 16.0% | 16.2% |
| Failure rate | 0.7% | 0.7% |

Table S1.5. Assay Limits in EU/ml. Not adjusted for dilutions.

| | **oFANG** | **nFANG** | **Q2** | **Luminex** |
|---|---|---|---|---|
| LOD | 31.74 | 27.14 | 0.0009 | 6.68 |
| LLOQ | 55.34 | 66.96 | 0.0049 | 2.99 |
| ULOQ | 2,511.89 | 55,526.77 | 20 | 1840.60 |

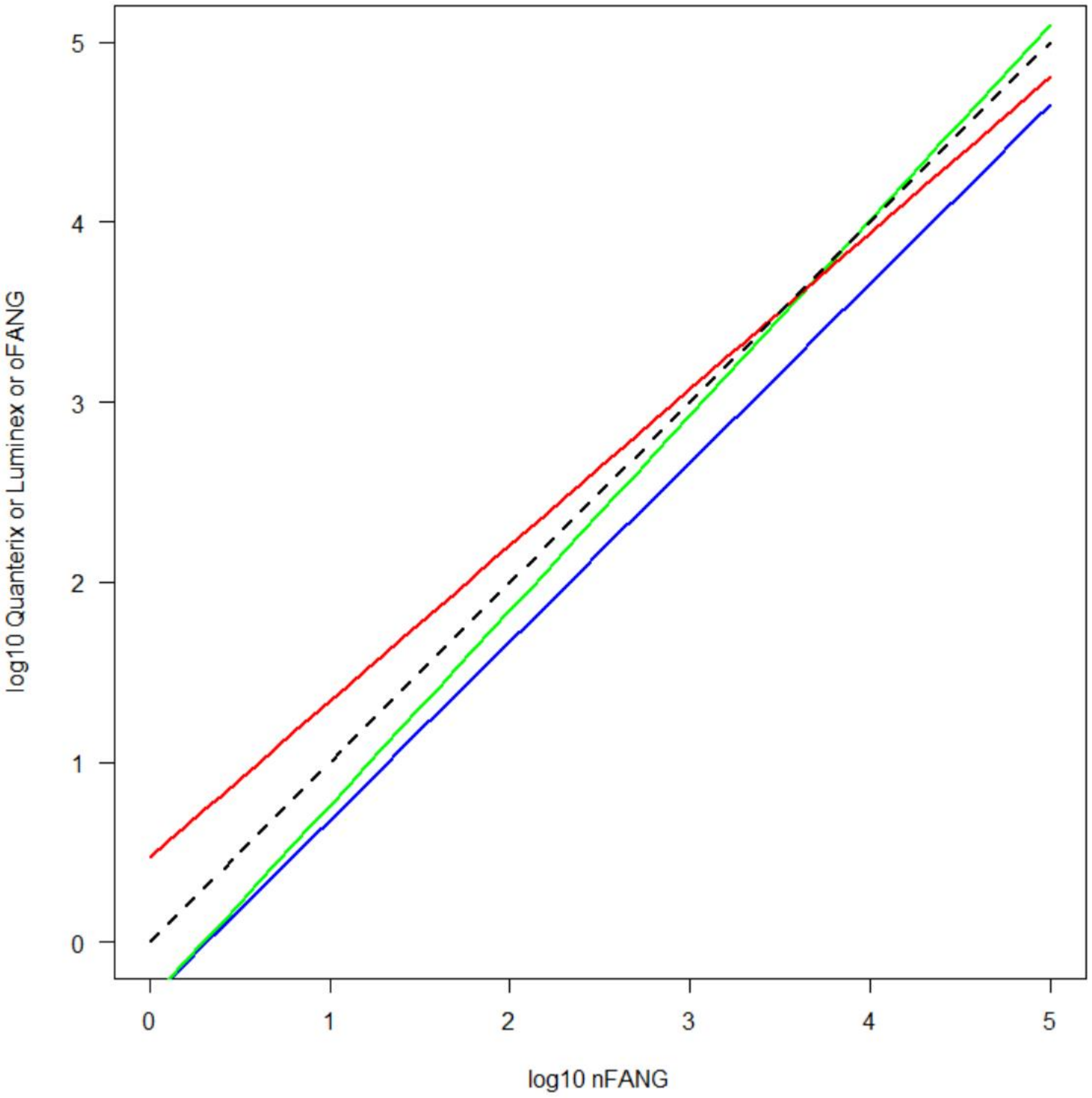

Figure S1.1. Deming regressions of oFANG (red), Luminex (green), and Q2 (blue) assays versus the nFANG assay. The black dashed line is a 45-degree reference line.

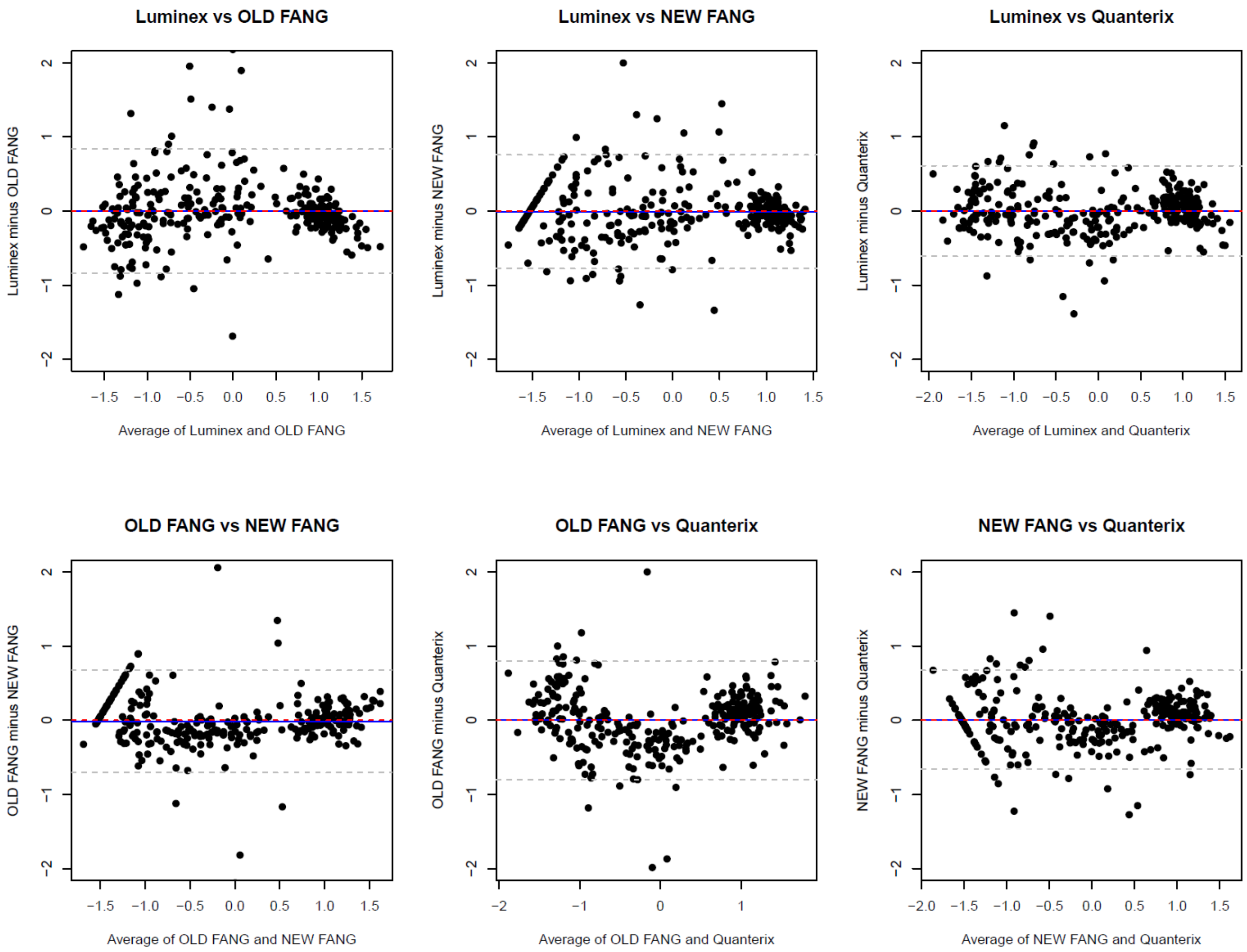

Figure S1.2. Bland-Altman plots for the four different assays (two-by-two comparisons).

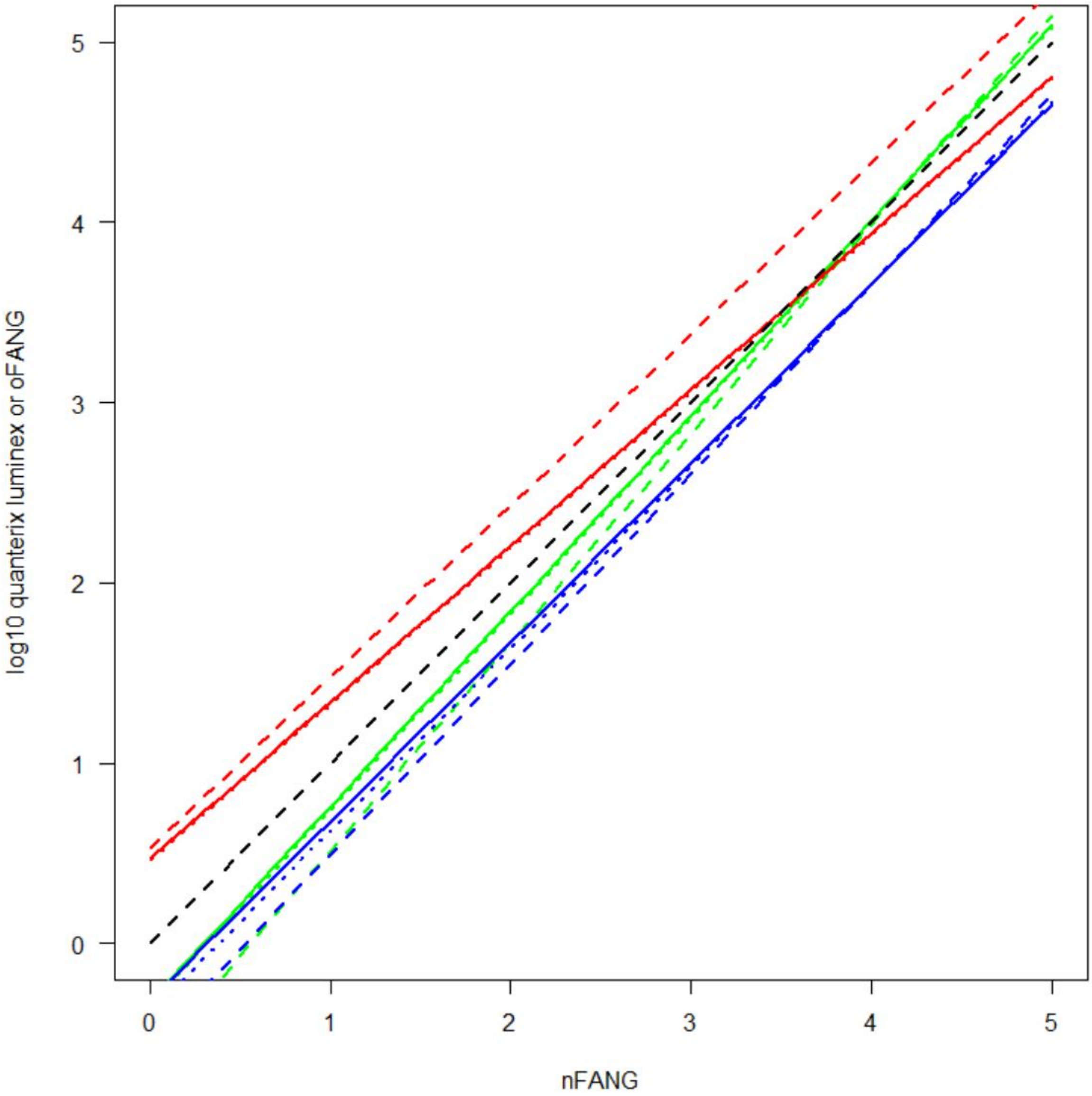

Figure S1.3. Visualization of the conversion equations from nFANG to the other assays: oFANG (red), Luminex (green), and Quanterix (blue) assays versus the nFANG assay under different methods of handling values below LOD (solid=LOD/2, dashed=remove, dotted=raw).

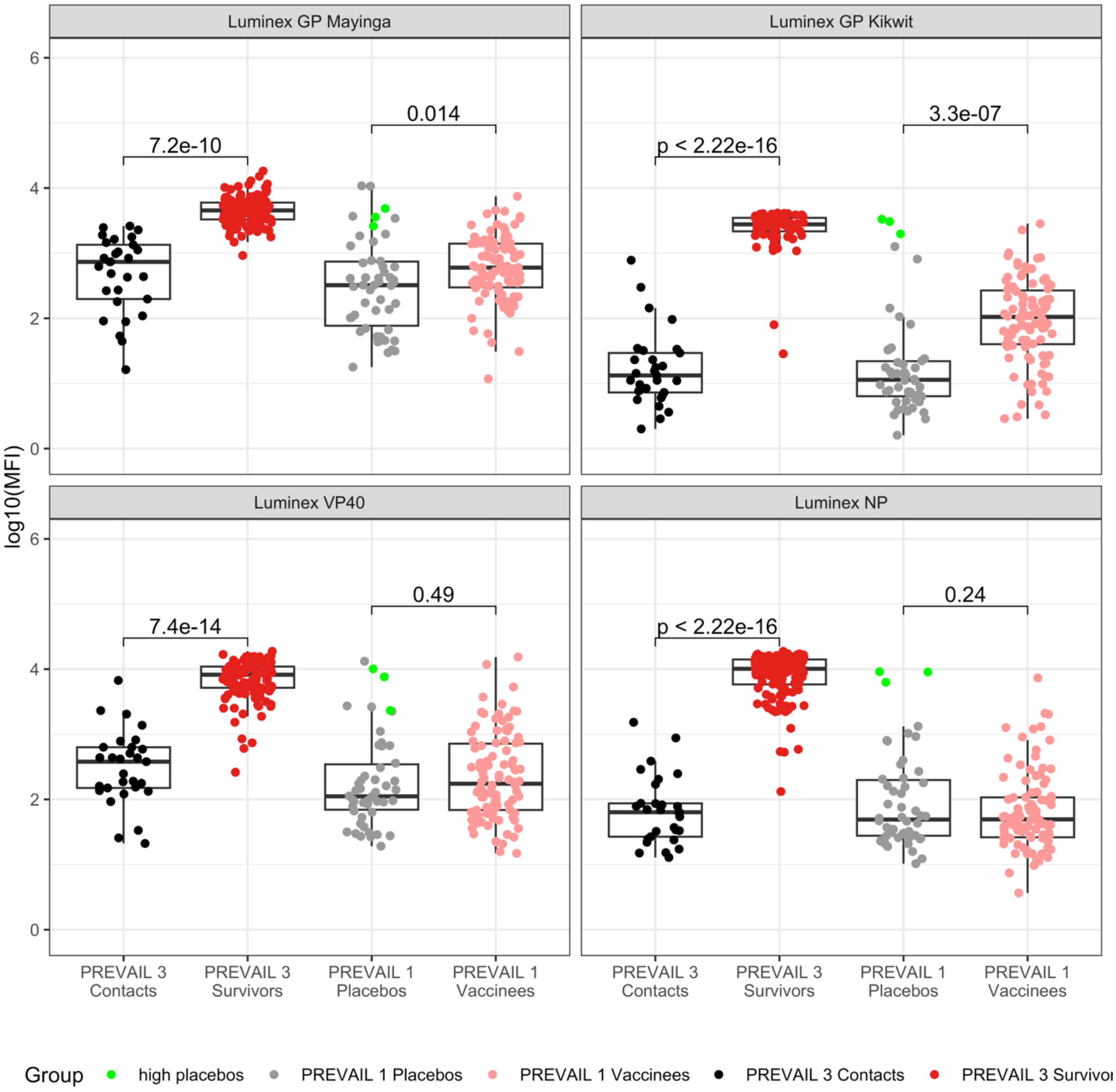

Figure S1.4. Dot plots of the four different targets of the Luminex assay for the four groups of participants; PREVAIL 3 contacts and survivors and PREVAIL 1 vaccine and placebo arms. Three placebo arm participants that read high are denoted by green dots.

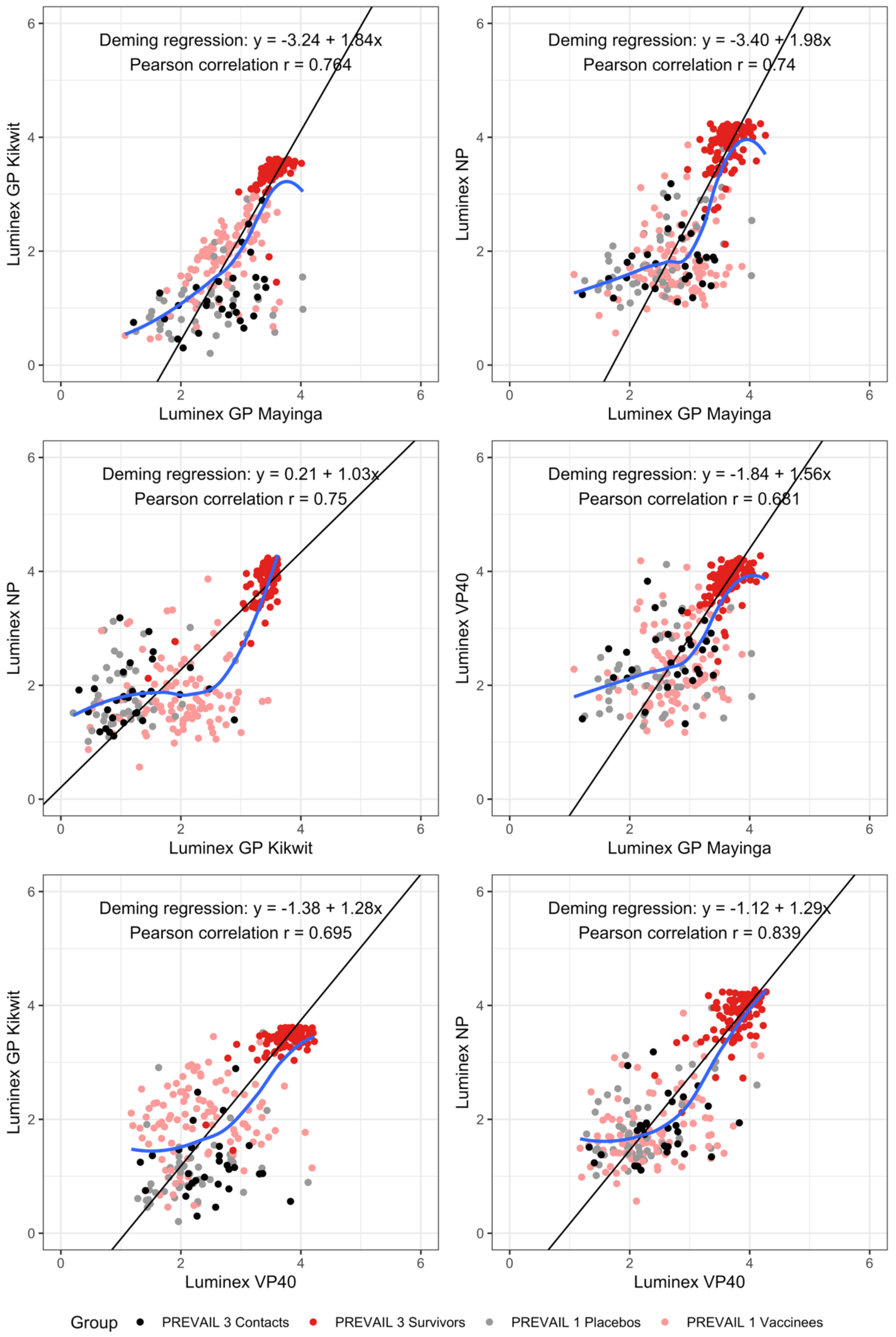

Figure S1.5. Pairwise scatterplots of the four different targets of the Luminex assay.

#### 2. ASSAY VALIDATION: LUMINEX

**Materials and methods**

**Panels sample and controls**

We used a panel of 85 human samples of known Ebola virus serostatus to validate our Luminex based multiplex immunoassay. All the human sample used in this work were anonymized with no way to link back to the 85 patients. The panel included 42 sera from the Virology department of the University Hospital of Montpellier were used as negative control and 43 sera coming from a cohort of Ebola virus disease survivor of 2013-2016 outbreak in Guinea were used as positive controls (Ayouba et al. JCM, 2017).

Three pooled serum control with anti-EBOV antibodies concentration information in ELISA Units by milliliter tested by ELISA Fang (Rudge et al, PlosOne, 2019) and two WHO standards (NIBSC) with anti-EBOV antibodies concentration information in international system unit were also used in this study (https://nibsc.org/products/brm_product_catalogue/detail_page.aspx?catid=15/220). To evaluate the assay robustness, we assessed intra- and inter-assay variabilities by testing known samples multiple times, previously tested on the FANG assay as Reference Standard, High Quality Control (HQC) and Low Quality Control (LQC) (Rudge et al. PlosOne, 2019).

Finally, two commercial monoclonal antibodies were used to validate our test: an anti-EBOV GP monoclonal antibody (LA63_CATB-B1092, CD, Shirley NY, USA) and an anti-Ebola surface glycoprotein (KZ52_Ab00960, Absolute Antibody, Wilton, UK).

**Luminex based multiplex immunoassay**

**Recombinant proteins**

We used as targets for our Luminex based multiplex immunoassay four commercially available recombinant proteins from different genomic region of orthoebolaviruses (nucleoprotein variant/isolate Makona-Kissidougo-C15 2014 (NP) (40443-V07E1 – Sinobiological (China)); viral protein 40 Makona-Kissidougo-C152014 (VP40) (40446-V07E – Sinobiological (China)); glycoprotein isolate Mayinga 1976 (GP-m) (40304-V08B1– Sinobiological (China)); and glycoprotein strain Kikwit 1995 (GP-k) (EBOVKW95-ENV – The Native Antigen Company, Oxford, UK). The proteins were purchased as lyophilized powders with purity above 90% as defined by SDS-PAGE. They were resuspended in solution and stored as per manufacturer's instruction until use.

**Proteins coupling to Luminex Beads**

We used our previously described protocol for coupling recombinant proteins to Luminex beads (Ayouba et al. JCM, 2017). Briefly, recombinant proteins (2μg/$10^6$ beads) were covalently coupled on carboxyl functionalized fluorescent magnetic beads (Luminex Corp., Austin, TX) with the BioPlex amine coupling kit (Bio-Rad Laboratories, Marnes-la-Coquette, France) according to the manufacturer's instructions. We blocked unreacted sites with blocking buffer from the amine coupling kit. Protein-coupled microsphere preparations were washed with PBS and stored in storage buffer from the amine coupling kit at 4˚C in the dark until use.

**Luminex based multiplex immunoassay for IgG antibody in serum**

Before use, recombinant protein-coupled beads are vortexed for 30 seconds and diluted to 2,000 beads/μL of assay buffer (Phosphate Buffered Saline (PBS) containing 0.75 mol/L NaCl, 1% (wt/vol) bovine serum albumin (Sigma Aldrich, Saint-Quentin Fallavier, France), 5% (vol/vol) heat-inactivated fetal bovine serum (Gibco-Invitrogen, Cergy Pontoise, France), and 0.2% (vol/vol) Tween-20 (Sigma-Aldrich). Tests are performed in 96-well flat-bottom chimney plates (Greiner bio one, Frickenhausen, Germany) and 50 μL of bead mixture is added to each well. Then, the supernatant is aspirated using an automatic plate washer (BioTek 405TS Microplate washer). Next, wells are incubated with 100 µL of plasma (diluted 1/200 in assay buffer) for 16 hours at 4˚C in the dark on a plate shaker at 400 rpm/min.

After five washings with 100 µL of assay buffer, 50 µL of biotin-labeled anti-human IgG is added (BD-Pharmingen, Le Pont De Claix, France) at a concentration of 4 µg/mL in each well and incubated for 30 minutes in the dark with continuous shaking at 400 rpm/min. Plates are washed five times as above, and 50 µL of streptavidin-R-phycoerythrin (Fisher Scientific/Life Technologies, Illkirch, France) at 4 µg/ml are added per well and incubated for 10 minutes while shaking at 400 rpm/min. The washer performs another five washes, then 150 µL of reading buffer (PBS 1X, 1% de BSA) is added per well to read antigen-antibody reactions on Intelliflex (Luminex Corp., Austin, TX, USA). At least 100 events are read for each bead set, and the results are expressed as median fluorescence intensity (MFI) per 100 beads.

**Assay performance determination**

To determine the performance, we determined for each of the four proteins (NP, VP40, GPm and GPk), sensitivity, specificity, and accuracy with ROC curve analysis with XLSTAT (Addinsoft, Paris, France) implemented in Microsoft Excel software.

To define the assay consistence, we determined assay variabilities by calculating intra- and inter-assay variability. We determined the limits of quantification for both Glycoproteins (GPm and GPk) by using the two monoclonal antibodies as samples. We used the Belysa software (Merck-Millipore) to determine the upper and lower limits of quantification of the assay.

#### Results

##### Sensitivity, specificity and accuracy

We used the panel of 85 samples (43 positive and 42 negative) of known Ebola virus to determine the intrinsic performance of our test.

The results are summarized in the Table S.2.1 below and Figure S.2.1 and show excellent performances for all the four antigens, with accuracies (Area Under the Curve, AUC) comprised between 98.7% and 100%.

**Table S2.1:** Sensitivity, specificity, accuracy, and cut-off values of the four antigens used in the Luminex assay.

| Antigens | Sensitivity (%) | Specificity (%) | Accuracy (%) | Cut-off (MFI) |
|---|---|---|---|---|
| **GPk** | 100 | 100 | 100 | 169 |
| **GPm** | 97.7 | 92.9 | 98.7 | 1081 |
| **NP** | 100 | 100 | 100 | 590 |
| **VP40** | 100 | 100 | 100 | 869 |

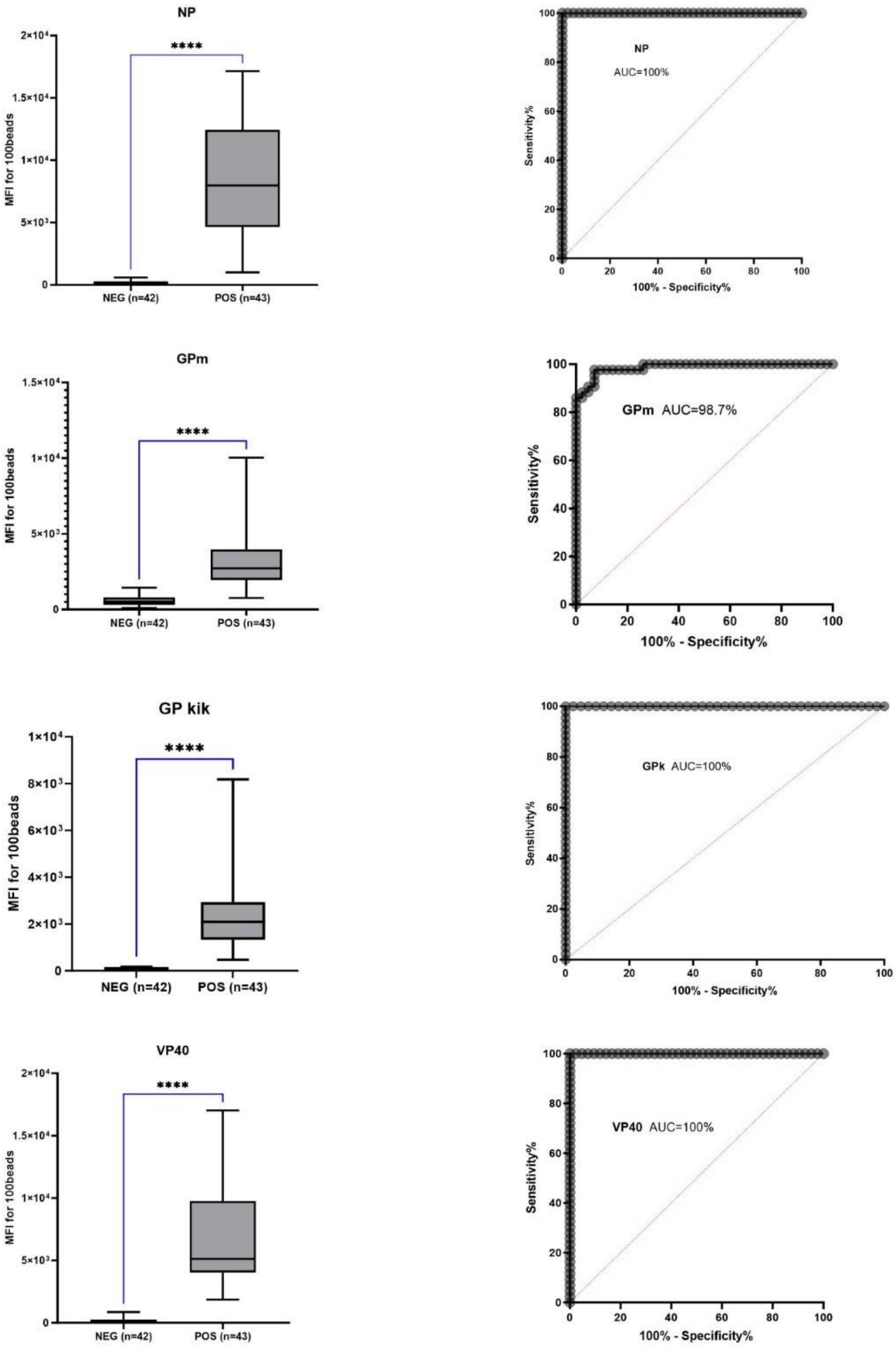

**<u>Figure S2.1:</u>** ROC curves of the four antigens used in the Luminex assay, together with boxplot comparison of negative and positive samples reactivities on the same four antigens. The statistical comparison was performed with the Mann-Whitney U-test. All p-values are highly significant ($<10^{-4}$). NP, Ebola virus nucleoprotein; GP, Ebola virus glycoprotein, VP40, Ebola virus 40kDa viral protein.

**Intra-assay variability**

To evaluate the robustness of our assay, we first assessed the intra-assay variability. To that end, we tested in the same run 12 replicates of each of the three references (i.e. reference Standard; HQC and LQC). From these 12 replicates, we calculated the coefficient of variation (CV) for each sample.

Results of this assessment are provided below, in percentage. It should be noted that for all the four antigens, the CV is below 20%.

**Table S2.2:** Intra-assay variability for the four antigens used, tested in 12 replicates within the same run.

| | Intra-assay variability (%) | | | |
|---|---|---|---|---|
| | NP | GPmay | VP40 | GPkikwit95 |
| Ref Standard | 11.38 | 10.26 | 12.69 | 13.51 |
| HQC | 12.86 | 11.62 | 14.40 | 12.02 |
| LQC | 18.76 | 9.90 | 13.25 | 11.72 |

**Inter-assay variability**
Next, we assessed the variability of the assay among plates ran on separate days by two different engineers. The same controls were used. For the reference standard, HQC (High Quality Control) and LQC (Low Quality Control), 19, 22, and 18 values were recorded, respectively. The results of the inter-assay variability are presented in Table S2.3 below and the showed that for all the four antigens the CVs are below 30%.

**Table S2.3:** inter-assay variability for the four antigens used, tested in 18-22 replicates in multiple runs.

| | Inter-assay variability (%) | | | |
|---|---|---|---|---|
| | NP | GPmay | VP40 | GPkikwit95 |
| Ref standard | 26.94 | 26.37 | 24.53 | 20.80 |
| HQC | 17.25 | 26.77 | 17.68 | 16.76 |
| LQC | 22.72 | 19.42 | 18.71 | 14.03 |

**Determination of detection limits of antibodies against EBOV GP kikwit95 and Mayinga76.**
To that end, we used two monoclonal antibodies raised against the two glycoproteins.
Figure S2.2 below illustrates such a titration curve for GPkikwit95 with the monoclonal antibody KZ52. The LLoQ is 112pg/ml, the MDD (Minimum Detectable Dose) is 89.26 pg/ml and the ULQ (Upper Limit of Quantification) is 10,000pg/ml.

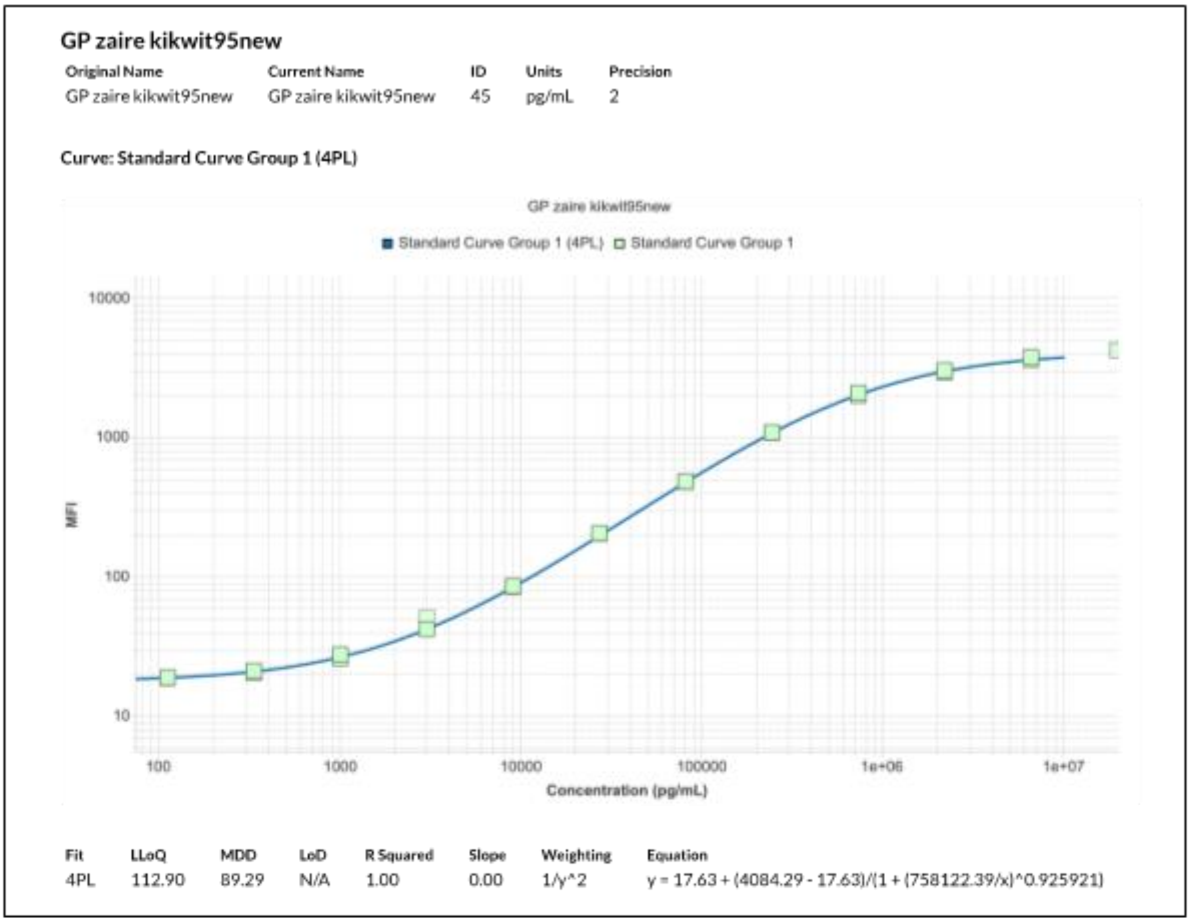

**Figure S2.2:** Titration curve of GPkikwit95 with the monoclonal antibody KZ52. Trend and limits for the MoAb LA63 against GPmayinga are similar.

#### 3. ASSAY VALIDATION: QUANTERIX

| | |
|---|---|
| **Test facility:** | Quanterix,<br>900 Middlesex Turnpike, Building 1, Billerica, MA 01821 |
| **Sponsor:** | NIAID/Leidos |
| Sponsor's Monitor: | Kevin Newell |

| | |
|---|---|
| **Dates:** | |
| Study initiation: | 19 Aug 2022 |
| Experimental start: | 06 Sep 2022 |
| Experimental completion: | 29 Sep 2022 |
| Final report: | 26 Aug 2024 |

| | |
|---|---|
| **Archive:** | Quanterix, 900 Middlesex Turnpike, Building 1, Billerica, MA 01821 |
| **Personnel involved in the study:** | |
| Study Responsible: | Scott Van Arsdell, PhD<br>Quanterix, 900 Middlesex Turnpike, Building 1, Billerica, MA 01821<br><br>Susan Yan, PhD<br>Quanterix, 900 Middlesex Turnpike, Building 1, Billerica, MA 01821 |

#### Table of figures

Page

**No table of figures entries found.**

**Table of tables**

**Page**

**Abbreviations**

| **Abbreviation** | **Definition** |
|---|---|
| % CV | Percent coefficient of variation |
| ALQ | Above Limit of Quantification |
| BAR | Biotinylated Antibody Reagent |
| BLD | Below Limit of Detection |
| BLQ | Below Limit of Quantification |
| CCD | Couple-Charged Device |
| EBOV | Ebola Virus |
| ELISA | Enzyme Linked Immunosorbent Assay |
| HQC | High Quality Control |
| HRP | Horseradish Peroxidases |
| IV | Intensity Value |
| LLOQ | Lower Limit of Quantification |
| LOD | Limit of Detection |
| LQC | Low Quality Control |
| mg | milligram |
| mL | milliliter |
| MRD | Minimum required dilution |
| MQC | Medium Quality Control |
| µL | Microliter |
| pg | picogram |
| QC | Quality Control |
| % RE | Percent relative error |
| SA-HRP | Streptavidin HRP |
| SD | Standard Deviation |
| SOP | Standard Operating Procedure |
| ULOQ | Upper Limit of Quantification |
| VHQC | Very High Quality Control |
| VLQC | Very Low Quality Control |

**Brief summary**

The SP-X platform is a multiplex sandwich enzyme-linked immunosorbent assay (ELISA) in a planar, plate-based array format, for the quantitative measurement of secreted proteins in serum, plasma, tissue culture supernatants and other matrices.  The SP-X platform was used in this project to develop and validate a serology assay for the detection of IgG antibodies to EBOV GP monomer protein in human serum.  Each well of the microplate is pre-spotted with GP monomer protein, which binds to anti-GP monomer antibodies in the standards, controls and samples added to the plate.  After unbound antibodies and serum components are washed away, a biotinylated anti-IgG detection antibody is added that binds to IgG antibodies bound to the GP monomer target protein. After washing away excess detection antibody, streptavidin-horseradish peroxidase (SA-HRP) is added.  The HRP enzyme conjugate subsequently reacts with the chemiluminescent substrate to produce a luminescent signal that is detected using the SP-X CCD imaging and analysis system. The amount of signal produced is proportional to the amount of anti-GP monomer antibody in the original standard or sample.  Customized SP-X software uses a five-parameter curve fit to back calculate unknowns using results extrapolated from the corresponding standard curve.

The objective of this study was to validate a planar serology immunoassay for the detection of IgG antibodies to EBOV GP monomer protein in human serum samples.  Validation studies included an assessment of standard curve performance, determination of the limit of quantification (LLOQ) with human serum samples, intra-assay and inter-assay precision with spiked normal human serum samples, dilutional linearity with human serum samples, spike and recovery with human serum samples, and a reference range study with serum samples from patients who have been vaccinated with the EBOV vaccine. The results of the study indicate that the GP monomer serology immunoassay is valid and suitable for clinical sample analysis.

### 1 Introduction

The SP-X platform is a multiplex sandwich enzyme-linked immunosorbent assay (ELISA) in a planar, plate-based array format, for the quantitative measurement of secreted proteins in serum, plasma, tissue culture supernatants and other matrices. The SP-X platform was used in this project to develop and validate a serology assay for the detection of IgG antibodies to EBOV GP monomer protein in human serum. Each well of the microplate is pre-spotted with GP monomer protein, which binds to anti-GP monomer antibodies in the standards, controls and samples added to the plate. After unbound antibodies and serum components are washed away, a biotinylated anti-IgG detection antibody is added that binds to IgG antibodies bound to the GP monomer target protein. After washing away excess detection antibody, streptavidin-horseradish peroxidase (SA-HRP) is added. The HRP enzyme conjugate subsequently reacts with the chemiluminescent substrate to produce a luminescent signal that is detected using the SP-X CCD imaging and analysis system. The amount of signal produced is proportional to the amount of anti-GP monomer antibody in the original standard or sample. Customized SP-X software uses a five-parameter curve fit to back calculate unknowns using results extrapolated from the corresponding standard curve.

The objective of this study was to validate a planar serology immunoassay for the detection of IgG antibodies to EBOV GP monomer protein in human serum samples. Validation studies included an assessment of standard curve performance, determination of the limit of quantification (LLOQ) with human serum samples, intra-assay and inter-assay precision with spiked normal human serum samples, dilutional linearity with human serum samples, spike and recovery with human serum samples, and a reference range study with serum samples from patients who have been vaccinated with the EBOV vaccine. The results of the study indicate that the GP monomer serology immunoassay is valid and suitable for clinical sample analysis.

### 2 Materials and methods

#### 2.1 Array

A planar serology immunoassay array was constructed for detection of the following target antibody:

EBOV GP monomer IgG

#### 2.2 Materials

Each assay consisted of a 96-well plate custom arrayed with GP monomer protein, a reference standard serum, sample diluent containing 0.1% sodium azide, biotinylated antibody reagent, SA-HRP reagent, stable peroxide solution, luminol enhancer solution, and wash buffer. The lot numbers and expiration dates for the critical assay components are presented in Table 3.1. Plates were read using the SP-X CCD camera imaging and analysis system. Images were analysed using SP-X software and data analysis was completed and summarized using Microsoft® Excel.

#### 2.3 Methods

Prior to running assays, all reagents were brought to room temperature. Plates were washed using the Quanterix plate washer SP-X program. Diluted samples, standards and controls were incubated for two hours on the arrayed plates. All incubations were performed at ambient temperature with shaking at 450 rpm. Plates were washed using the Quanterix plate washer SP-X program before adding the biotinylated detection antibody to each well. After incubating with detection antibody for 30 minutes, plates were washed and incubated for 30 minutes with streptavidin-horseradish peroxidase. Plates were again washed before adding the chemiluminescent substrate. The plates were then imaged using the SP-X imaging system and data was analysed using the SP-X software. Concentrations of all unknown samples were back calculated using results extrapolated from the corresponding standard curve. A five-parameter curve fit was used for all standard curves.

##### 2.4 Preparation of controls

Control samples corresponding to very high, high, medium, low, and very low concentration levels (VHQC, HQC, MQC, LQC, VLQC) were created for the GP monomer serology assay. A serum sample with high levels of anti-EBOV GP monomer antibodies (lot number PREP0001, received from NIAID) was spiked into two normal human serum samples (BRH233209 and HMN403957) at five concentration levels (VHQC, HQC, MQC, LQC, VLQC) for each sample to create ten individual control samples for the validation. Two intermediate controls for each concentration level (pre-VHQC-1, pre-VHQC-2, pre-HQC-1, pre-HQC-2, pre-MQC-1, pre-MQC-2, pre-LQC-1, pre-LQC2, pre-VLQC-1 and pre-VLQC-2) were manufactured, aliquoted into individual vials and stored at -80°C. The intermediate controls were diluted to the final target concentrations for VHQC-1, VHQC-2, HQC-1, HQC-2, MQC-1, MQC-2, LQC-1, LQC2, VLQC-1 and VLQC-2 on the day of the assay. The lot numbers and expiration dates of the reagents used to manufacture quality control samples are presented in Table 3.2. The final target concentrations for the quality control samples are presented in Table 3.

#### 3 Archiving procedures

Data and the report will be stored at Quanterix, 900 Middlesex Turnpike, Building 1, Billerica, MA 01821. Electronic data are backed up daily. All documents and data related to the assay validations and sample testing service will be stored for five years by Quanterix.

#### 4 Results

##### 4.1 Analytical Run Summary

Validation of the EBOV GP monomer planar serology immunoassay was performed over 12 analytical runs presented in Table 3.4. Twelve analytical runs were accepted.

##### 4.2 Standard Curve Performance

The standard curve assay range for the EBOV GP monomer serology assay was determined during development and was selected to optimize sensitivity and linearity. Standard curves were run in assay Diluent 22. A minimum of six standard curves were evaluated over the course of the validation. Seven non-zero calibrators in triplicate within the anticipated range were included for a concentration-response relationship fit using the five-parameter logistic function. The standard curve was generated by serially diluting standard calibrator 1 with four-fold intervals.
Standard curve performance is acceptable if the CV of the mean signal (integrated density value) for standards 1-5 is < 15%, the CV of the mean signal (integrated density value) for standards 6 and 7 is <20%, the percent recovery for the calculated back fit concentration for standards 1-6 is 80 – 120% and the percent recovery for the calculated back fit concentration for standard 7 is 75 – 125%.
The following equations were used to calculate % CV and % RE.
% CV = standard deviation ÷ mean signal of duplicate wells x 100
% Recovery = measured concentration ÷ nominal concentration x 100
% RE = (measured concentration – nominal concentration) ÷ nominal concentration x 100
The upper and lower limits of quantification (ULOQ and LLOQ) for each analyte was established based on the accuracy and repeatability of the high and low reference standards. The limit of

detection (LOD) was calculated by adding 2.55 standard deviations of signal to the mean signal of the replicates of the zero standards and determining the resulting concentration from the standard curve. The Lower Limit of Quantification (LLOQ) of the assay was defined as the lowest calibration standard with a back-calculated concentration CV of less than 20% and back-calculated concentration relative error of less than 25%. The Upper Limit of Quantification (ULOQ) of the assay was defined as the highest calibration standard with a back-calculated concentration of less than 20% and back-calculated concentration relative error of less than 20%.
A typical standard curve for the EBOV GP monomer serology assay is presented in Table 3.5. The analytical ULOQ, LLOQ and LOD for the EBOV GP monomer serology assay is presented in Table 3.6. The LOD shown in Table 3.6 is the average LOD value from 10 calibration curves. The backfit concentrations of each calibrator for each EBOV GP monomer assay run of the validation (n = 12 runs) is presented in Table 3.7. The mean backfit value, standard deviation, % CV, % nominal and % RE are also presented.

#### 4.3 Limit of Quantification with Sample Matrix

The LLOQ of the EBOV GP monomer serology assay was also determined with sample matrix. Four serum samples from NIAID with levels of anti-EBOV GP monomer antibodies that are above the LLOQ determined with the calibration standards were used for this analysis. Each sample was diluted 1:4,000, 1:8,000, 1:16,000, 1:32,000, 1:64,000, 1:128,000, 1:256,000 and 1:512,000 in sample diluent and then run on the assay. The LLOQ for sample matrix was defined as the lowest analyte concentration that can quantified in serum with acceptable precision (+/- 25%). The concentration data for each sample replicate for each sample dilution is presented in Table 3.8. The LLOQ values determined with sample matrix for each sample are also shown in Table 3.8. The average LLOQ value determined with sample matrix was 0.002 EU/ml.

#### 4.4 Intra-Assay Precision

Control samples VHQC-1, VHQC-2, HQC-1, HQC-2, MQC-1, MQC-2, LQC-1, LQC2, VLQC-1 and VLQC-2 were created as described in Section 2.4. Intra-assay precision was evaluated by running 16 replicates each of VHQC-1, HQC-1, MQC-1, LQC-1 and VLQC-1 on one plate and 16 replicates each of VHQC-2, HQC-2, MQC-2, LQC-2 and VLQC-2 on a second plate. The % CV of back-calculated protein concentration was calculated for each control sample. The intra-assay precision was considered acceptable if % CV was ≤25%. Intra-assay precision data is presented in Table 3.9 and Table 3.10.

#### 4.5 Inter-Assay Precision

Inter-assay precision was evaluated by running duplicates of VHQC-1, VHQC-2, HQC-1, HQC-2, MQC-1, MQC-2, LQC-1, LQC2, VLQC-1 and VLQC-2 on at least four assay plates on four separate days, using at least two operators and calculating % CV of back-calculated protein concentration for each control sample. The inter-assay precision was considered acceptable if % CV was ≤25%. Inter-assay precision data is presented in Table 3.11 and Table 3.12.

#### 4.6 Dilutional Linearity/Parallelism

Dilution and recovery experiments were performed to assess the dilutional linearity of the EBOV GP monomer serology assay. Five serum samples from NIAID with high levels of anti-EBOV GP monomer antibodies were used for this analysis. Each sample was diluted 1:2,000, 1:4,000, 1:8,000, 1:16,000 and 1:32,000 in sample diluent and then run on the assay. The measured concentration of each sample dilution was multiplied by the dilution factor and compared to the concentration of the 1:4,000 dilution. Recoveries of each of the dilutions was calculated as a percent recovery of protein relative to the 1:4,000 dilution. The 1:8,000 sample dilution was also used as a reference for calculating % recovery. Dilutional linearity is considered acceptable if recovery is ±30% of the reference concentration value. The antibody concentrations of the diluted samples are shown in Table 3.13 and the percent recoveries for each sample are shown in Table 3.14.

##### 4.7 Spike and Recovery

Spike and recovery experiments were performed to assess serum matrix effects and the accuracy of the method in serum samples. Serum samples received from NIAID were spiked with a serum sample containing high levels of anti-EBOV GP monomer antibodies (lot number PREP0001, received from NIAID). Ten individual serum samples were diluted at 1:2000 with sample diluent and spiked at very high, medium, low and very low concentrations of anti-EBOV GP monomer antibodies spanning the calibration range of the assay. The very high spike was targeted to fall between standards 1 and 2, the high spike between standards 2 and 3, the medium spike between standards 3 and 4, the low spike between standards 4 and 5 and the very low spike between standards 5 and 6. Each serum sample was also assayed without a spike to measure endogenous levels for use in recovery calculations. The final sample dilution after the spike solution was added was 1:4,000. The measured value from the un-spiked matrix-matched sample is subtracted from the measured values of each of the spiked samples. The net value was compared to the measured value of the five concentrations of protein spiked into the control diluent. The sample matrix effect is represented as a percent recovery of expected protein concentration. Percent recovery was calculated using the following equation:
% Recovery = (spiked sample concentration – endogenous) ÷ control spike concentration x 100
Spike and recovery is considered acceptable if the recovery of each analyte in ≥ 5/7 samples is within ±30% of the control spike concentration. The measured concentrations of anti-EBOV antibodies in spiked samples are presented in Table 3.15 and Table 3.17. The percent recovery for each of the spiked samples is presented in Table 3.16 and Table 3.18.

##### 4.8 Serum Sample Testing

Sixty serum samples from vaccinated individuals received from the IRF were analysed with the EBOV GP monomer serology assay. Samples were diluted 1:4,000 with sample diluent and then run on the assay. The measured concentration of anti-EBOV GP monomer antibodies for each sample is presented in Table 3.19 and Table 3.20.

#### 5 Discussion

##### 5.1 Standard Curve Performance

All standard curves for the validation assay runs met the specifications for precision and accuracy described in Section 4.2. The backfit concentrations of each calibrator for each assay run (n = 12) are presented in Table 3.7. The mean backfit value, standard deviation, % CV, % nominal, and % RE are acceptable for all calibration curves run for the validation study.

##### 5.2 Limit of Quantification with Sample Matrix

The LLOQ for sample matrix was defined as the lowest analyte concentration that can be quantified in serum with acceptable precision (+/- 25%). The LLOQ values determined with sample matrix for the EBOV GP monomer serology assay are shown in Table 3.8. The average LLOQ value determined with sample matrix was 0.002 EU/ml.

##### 5.3 Intra-Assay Precision

Intra-assay precision data is presented in Table 3.9 and Table 3.10. The intra-assay precision is considered acceptable as the % CV for all quality control samples for all assays was ≤25%.

##### 5.4 Inter-Assay Precision

Inter-assay precision data is presented in Table 3.11 and Table 3.12. The inter-assay precision is considered acceptable as the % CV for all quality control samples for all assays was ≤25%.

##### 5.5 Dilutional Linearity/Parallelism

Dilution and recovery experiments were performed to assess the dilutional linearity of the EBOV GP monomer serology assay. The antibody concentrations of the diluted samples are shown in Table 3.13 and the percent recoveries for each of the five samples are shown in Table 3.14. Dilutional linearity is considered acceptable if recovery of at least 2/3 (67%) of the dilution comparisons for 80% of the individual serum samples is within 70-130%. Dilutional linearity was acceptable for the five serum samples analysed whether the 1:4,000 or 1:8,000 dilution was used as the reference concentration (Table 3.14).

##### 5.6 Spike and Recovery

The results for spike and recovery analysis are shown in Table 3.15-Table 3.18. The measured concentrations of anti-EBOV antibodies in spiked samples are presented in Table 3.15 and Table 3.17. The percent recovery for each of the spiked samples is presented in Table 3.16 and Table 3.18. Spike recovery is considered acceptable if recovery of 4/5 (80%) of the spike levels for 80% of the individual samples is within 70-130% of the control diluent spike. Spike recovery was acceptable for all ten serum samples at all five spike levels.

##### 5.7 Serum Sample Testing

Sixty serum samples from vaccinated individuals received from the IRF were analysed with the EBOV GP monomer serology assay. The measured concentrations of anti-EBOV GP monomer antibodies for each sample are presented in Table 3.19 and Table 3.20. The measured concentration of anti-EBOV GP monomer antibodies was within the quantitative range of the assay for 87% of the samples. Eight samples were below the functional LLOQ (19.6 EU/ml). The % CV for sample replicates was <25% for all samples tested.

#### 6 Deviations

There were no deviations for this study.

#### 7 Conclusion

The validation was approved, and the assays can be used for clinical sample analysis.

### Figures and Tables

#### Table 3.1: Assay Components

| Component | Lot Number | Expiration Date |
|---|---|---|
| GP Monomer Plate | 063022P | 30-Jun-2027 |
| Biotinylated Detection Antibody Reagent | 080122SY | 1-Aug-2025 |
| Reference Standard Serum | BMIZAIRE 108 | 30-Sep-2023 |
| Diluent 22 | 319097 | 31-Oct-2022 |
| Diluent 22 | 319115 | 7-Nov-2022 |
| Streptavidin HRP | 319747 | 5-May-2025 |

#### Table 3.2: Reagents for the Manufacture of Quality Control Samples

| Reagent | Lot Number | Expiration Date |
|---|---|---|
| High Titer Serum | PREP0001 | 2-Aug-2024 |
| Normal Human Serum | BRH233209 | 1-Sep-2023 |
| Normal Human Serum | HMN403957 | 1-Sep-2023 |

#### Table 3.3: Target Concentrations for Validation Quality Control Samples

| Sample | Target Concentration (EU/ml) |
|---|---|
| VHQC | 10 |
| HQC | 2.5 |
| MQC | 0.625 |
| LQC | 0.156 |
| VLQC | 0.039 |

#### Table 3.4: Analytical Run Summary for the EBOV GP Monomer Serology Assay Validation

| Assay Run | Date | Plate Identity | Status | Tables |
|---|---|---|---|---|
| 1 | 6-Sep-2022 | serum dil linearity ebovgpmono 063022p-012 as0073 220906 151036.sdf | Accepted | 7,11,12,13,14 |
| 2 | 7-Sep-2022 | plate1 loq ebovgpmono 063022p-013 as0073 220907 150656.sdf | Accepted | 7,8,11,12 |
| 3 | 7-Sep-2022 | plate2 loq ebovgpmono 063022p-014 as0073 220907 151300.sdf | Accepted | 7,8,11,12 |
| 4 | 8-Sep-2022 | spk-rvy1 ebovgpmono 063022p-015 as0073 220908 144235.sdf | Accepted | 7,11,12,15,16 |
| 5 | 9-Sep-2022 | spk-rvy2 ebovgpmono 063022p-016 as0073 220909 151958.sdf | Accepted | 7,11,12,15,16,17,18 |
| 6 | 9-Sep-2022 | spk-rvy3 ebovgpmono 063022p-017 as0073 220909 152629.sdf | Accepted | 7,11,12,17,18 |
| 7 | 12-Sep-2022 | plate1 intra-precision ebovgpmono 063022p-018 as0073 220912 145455.sdf | Accepted | 7,9,11,12 |
| 8 | 12-Sep-2022 | plate2 intra-precision ebovgpmono 063022p-019 as0073 220912 150142.sdf | Accepted | 7,10 |
| 9 | 13-Sep-2022 | spk-rvy4 ebovgpmono 063022p-020 as0073 220913 160135.sdf | Accepted | 7,11,12,17,18 |
| 10 | 27-Sep-2022 | ebovgp az 1 1 ebovgpmono 063022p-023 as0073 220927 151153.sdf | Accepted | 5,7,11,12 |
| 11 | 29-Sep-2022 | plate3 irf13-45 serum samples ebovgpmono 063022p-025 as0073 220929 173534.sdf | Accepted | 7,19,20 |
| 12 | 29-Sep-2022 | plate4 irf61-95 serum samples ebovgpmono 063022p-026 as0073 220929 174221.sdf | Accepted | 7,19,20 |

#### Table 3.5: Typical Standard Curve for the EBOV GP Monomer Serology Assay

| Calibrator | GP Monomer (EU/ml) | Average Signal (IV) | SD | % CV | % RE |
|---|---|---|---|---|---|
| 1 | 20.0 | 4638347 | 153965 | 3.3 | 4.4 |
| 2 | 5.00 | 1985204 | 167534 | 8.4 | 1.1 |
| 3 | 1.25 | 580815 | 9260 | 1.6 | 7.3 |
| 4 | 0.313 | 164173 | 5689 | 3.5 | 5.1 |
| 5 | 0.078 | 39289 | 1175 | 3.0 | 8 |
| 6 | 0.0195 | 8561 | 976 | 11.4 | 1.5 |
| 7 | 0.0049 | 1733 | 204 | 11.8 | 20.7 |
| 8 | 0.0000 | 331 | 34 | 10.2 | NA |

**Table 3.6: Analytical ULOQ, LLOQ, and LOD for the EBOV GP Monomer Serology Assay**

| GP Monomer | EU/ml |
|---|---|
| ULOQ | 20.0 |
| LLOQ | 0.0049 |
| LOD | 0.0009 |

**Table 3.7: Standard Curve Intermediate Precision for the EBOV GP Monomer Serology Assay**

| Plate | Assay Run | Assay Date | Std1 20.0 EU/ml | Std2 5.00 EU/ml | Std3 1.25 EU/ml | Std4 0.313 EU/ml | Std5 0.078 EU/ml | Std6 0.020 EU/ml | Std7 0.005 EU/ml |
|---|---|---|---|---|---|---|---|---|---|
| serum dil linearity ebovgpmono 063022p-012 as0073 220906 151036.sdf | 1 | 6-Sep-2022 | 20.5 | 4.64 | 1.29 | 0.327 | 0.085 | 0.018 | 0.004 |
| plate1 loq ebovgpmono 063022p-013 as0073 220907 150656.sdf | 2 | 7-Sep-2022 | 20.0 | 4.59 | 1.25 | 0.338 | 0.082 | 0.019 | 0.004 |
| plate2 loq ebovgpmono 063022p-014 as0073 220907 151300.sdf | 3 | 7-Sep-2022 | 20.4 | 4.60 | 1.27 | 0.319 | 0.084 | 0.020 | 0.004 |
| spk-rvy1 ebovgpmono 063022p-015 as0073 220908 144235.sdf | 4 | 8-Sep-2022 | 19.9 | 4.83 | 1.26 | 0.317 | 0.081 | 0.016 | * |
| spk-rvy2 ebovgpmono 063022p-016 as0073 220909 151958.sdf | 5 | 9-Sep-2022 | 20.4 | 4.79 | 1.27 | 0.340 | 0.083 | 0.018 | 0.004 |
| spk-rvy3 ebovgpmono 063022p-017 as0073 220909 152629.sdf | 6 | 9-Sep-2022 | 20.3 | 5.02 | 1.29 | 0.314 | 0.081 | 0.019 | 0.004 |
| plate1 intra-precision ebovgpmono 063022p-018 as0073 220912 145455.sdf | 7 | 12-Sep-2022 | 20.1 | 4.76 | 1.29 | 0.318 | 0.081 | 0.019 | 0.004 |
| plate2 intra-precision ebovgpmono 063022p-019 as0073 220912 150142.sdf | 8 | 12-Sep-2022 | 21.3 | 4.77 | 1.25 | 0.316 | 0.083 | 0.020 | 0.004 |
| spk-rvy4 ebovgpmono 063022p-020 as0073 220913 160135.sdf | 9 | 13-Sep-2022 | 21.2 | 4.76 | 1.20 | 0.303 | 0.082 | 0.020 | 0.004 |
| ebovgp az 1 1 ebovgpmono 063022p-023 as0073 220927 151153.sdf | 10 | 27-Sep-2022 | 20.9 | 4.94 | 1.16 | 0.329 | 0.084 | 0.020 | 0.004 |
| plate3 irf13-45 serum samples ebovgpmono 063022p-025 as0073 220929 173534.sdf | 11 | 29-Sep-2022 | 21.6 | 4.76 | 1.19 | 0.321 | 0.083 | 0.023 | 0.004 |
| plate4 irf61-95 serum samples ebovgpmono 063022p-026 as0073 220929 174221.sdf | 12 | 29-Sep-2022 | 19.4 | 4.99 | 1.16 | 0.342 | 0.084 | 0.021 | 0.004 |
| Mean | | | 20.5 | 4.79 | 1.24 | 0.324 | 0.083 | 0.019 | 0.004 |
| SD | | | 0.6 | 0.14 | 0.05 | 0.012 | 0.001 | 0.002 | 0.0003 |
| % CV | | | 3.1 | 2.95 | 4.09 | 3.6 | 1.8 | 8.4 | 7.0 |
| % Nominal | | | 102.5 | 95.8 | 99.1 | 103.6 | 105.9 | 99.8 | 80.3 |
| % RE | | | 2.5 | -4.2 | -0.9 | 3.6 | 5.9 | -0.2 | -19.7 |
| N | | | 12 | 12 | 12 | 12 | 12 | 12 | 11 |

*Cell without data indicates calibration point removed which did not meet acceptance criteria.

**Table 3.8: Determination of Lower Limit of Quantification with Sample Matrix (EBOV GP Monomer)***

| Sample | Dilution Factor | Replicate 1 (EU/ml) | Replicate 2 (EU/ml) | Replicate 3 (EU/ml) | Average (EU/ml) | SD | %CV | LLOQ |
|---|---|---|---|---|---|---|---|---|
| PREVAIL 9 | 4,000 | 0.108 | 0.108 | 0.101 | 0.105 | 0.004 | 3.8 | 0.001 |
| | 8,000 | 0.049 | 0.049 | 0.048 | 0.048 | 0.0005 | 1.0 | |
| | 16,000 | 0.027 | 0.024 | 0.022 | 0.024 | 0.002 | 9.3 | |
| | 32,000 | 0.010 | 0.011 | 0.010 | 0.010 | 0.000 | 2.9 | |
| | 64,000 | 0.005 | **0.004** | 0.006 | **0.005** | 0.001 | 19.3 | |
| | 128,000 | **0.002** | **0.002** | **0.003** | **0.003** | 0.0001 | 5.5 | |
| | 256,000 | **0.002** | **0.002** | **0.002** | **0.002** | 0.0002 | 10.7 | |
| | 512,000 | **0.001** | **0.001** | **0.002** | **0.001** | 0.0001 | 10.6 | |
| PREVAIL 27 | 4,000 | 0.253 | 0.362 | 0.324 | 0.313 | 0.055 | 17.6 | 0.002 |
| | 8,000 | 0.143 | 0.135 | 0.143 | 0.140 | 0.005 | 3.5 | |
| | 16,000 | 0.056 | 0.067 | 0.072 | 0.065 | 0.008 | 12.9 | |
| | 32,000 | 0.028 | 0.032 | 0.033 | 0.031 | 0.002 | 7.5 | |
| | 64,000 | 0.013 | 0.013 | 0.014 | 0.013 | 0.001 | 5.4 | |
| | 128,000 | 0.008 | 0.007 | 0.007 | 0.007 | 0.001 | 10.1 | |
| | 256,000 | **0.004** | **0.004** | **0.003** | **0.004** | 0.0005 | 13.7 | |
| | 512,000 | **0.002** | **0.002** | **0.002** | **0.002** | 0.0000 | 1.1 | |
| PREVAIL 79 | 4,000 | 0.267 | 0.271 | 0.278 | 0.272 | 0.006 | 2.1 | 0.002 |
| | 8,000 | 0.131 | 0.130 | 0.131 | 0.131 | 0.001 | 0.5 | |
| | 16,000 | 0.067 | 0.065 | 0.065 | 0.066 | 0.001 | 1.9 | |
| | 32,000 | 0.031 | 0.032 | 0.030 | 0.031 | 0.001 | 2.9 | |
| | 64,000 | 0.015 | 0.014 | 0.014 | 0.014 | 0.001 | 4.2 | |
| | 128,000 | 0.007 | 0.007 | 0.007 | 0.007 | 0.0004 | 6.0 | |
| | 256,000 | **0.003** | **0.003** | **0.003** | **0.003** | 0.0003 | 9.8 | |
| | 512,000 | **0.002** | **0.002** | **0.002** | **0.002** | 0.0001 | 5.6 | |
| PREVAIL 91 | 4,000 | 0.290 | 0.301 | 0.297 | 0.296 | 0.005 | 1.8 | 0.002 |
| | 8,000 | 0.146 | 0.144 | 0.142 | 0.144 | 0.002 | 1.4 | |
| | 16,000 | 0.072 | 0.069 | 0.072 | 0.071 | 0.002 | 2.2 | |

| | | | | | | | |
|---|---|---|---|---|---|---|---|
| | 32,000 | 0.035 | 0.034 | 0.034 | 0.035 | 0.0004 | 1.0 |
| | 64,000 | 0.015 | 0.015 | 0.016 | 0.015 | 0.0003 | 1.7 |
| | 128,000 | 0.008 | 0.007 | 0.007 | 0.007 | 0.0003 | 3.9 |
| | 256,000 | **0.004** | **0.003** | **0.004** | **0.004** | 0.0004 | 9.9 |
| | 512,000 | **0.002** | **0.002** | **0.002** | **0.002** | 0.0001 | 4.5 |

*Bold values are below lowest calibration standard

**Table 3.9: Intra-assay Precision Data for the EBOV GP Monomer Serology Assay (Plate 1)**

| Replicate | VHQC-1 (EU/ml) | HQC-1 (EU/ml) | MQC-1 (EU/ml) | LQC-1 (EU/ml) | VLQC-1 (EU/ml) |
|---|---|---|---|---|---|
| 1 | 9.03 | 2.09 | 0.517 | 0.163 | 0.069 |
| 2 | 8.25 | 1.91 | 0.514 | 0.156 | 0.071 |
| 3 | 8.48 | 1.92 | 0.492 | 0.149 | 0.074 |
| 4 | 8.60 | 1.98 | 0.491 | 0.154 | 0.071 |
| 5 | 8.33 | 2.00 | 0.476 | 0.150 | 0.073 |
| 6 | 8.93 | 1.83 | 0.554 | 0.157 | 0.074 |
| 7 | 8.45 | 2.10 | 0.560 | 0.155 | 0.069 |
| 8 | 9.24 | 2.12 | 0.562 | 0.170 | 0.080 |
| 9 | 8.57 | 2.17 | 0.564 | 0.155 | 0.076 |
| 10 | 7.79 | 1.95 | 0.507 | 0.149 | 0.089 |
| 11 | 7.89 | 1.95 | 0.484 | 0.142 | 0.073 |
| 12 | 7.49 | 1.87 | 0.507 | 0.152 | 0.072 |
| 13 | 8.49 | 1.91 | 0.482 | 0.160 | 0.084 |
| 14 | 7.83 | 2.00 | 0.489 | 0.156 | 0.081 |
| 15 | 9.07 | 2.00 | 0.494 | 0.144 | 0.080 |
| 16 | 8.95 | 2.21 | 0.562 | 0.145 | 0.079 |
| Average | 8.46 | 2.00 | 0.516 | 0.154 | 0.076 |
| SD | 0.514 | 0.109 | 0.033 | 0.007 | 0.006 |
| %CV | 6.1 | 5.5 | 6.4 | 4.7 | 7.4 |

**Table 3.10: Intra-assay Precision Data for the EBOV GP Monomer Serology Assay (Plate 2)**

| Replicate | VHQC-2 (EU/ml) | HQC-2 (EU/ml) | MQC-2 (EU/ml) | LQC-2 (EU/ml) | VLQC-2 (EU/ml) |
|---|---|---|---|---|---|
| 1 | 8.40 | 1.93 | 0.456 | 0.143 | 0.041 |
| 2 | 7.34 | 1.65 | 0.465 | 0.126 | 0.041 |
| 3 | 7.71 | 1.69 | 0.456 | 0.133 | 0.045 |
| 4 | 7.11 | 1.73 | 0.448 | 0.140 | 0.044 |
| 5 | 7.27 | 1.70 | 0.437 | 0.131 | 0.042 |
| 6 | 7.83 | 1.58 | 0.516 | 0.134 | 0.042 |
| 7 | 7.64 | 1.78 | 0.518 | 0.136 | 0.039 |
| 8 | 8.19 | 1.98 | 0.559 | 0.146 | 0.048 |
| 9 | 7.68 | 1.88 | 0.524 | 0.130 | 0.042 |
| 10 | 6.66 | 1.70 | 0.490 | 0.123 | 0.050 |
| 11 | 6.77 | 1.59 | 0.454 | 0.132 | 0.044 |
| 12 | 6.47 | 1.59 | 0.461 | 0.138 | 0.044 |
| 13 | 6.59 | 1.60 | 0.427 | 0.136 | 0.044 |
| 14 | 6.67 | 1.73 | 0.462 | 0.132 | 0.047 |
| 15 | 8.53 | 1.81 | 0.479 | 0.124 | 0.043 |
| 16 | 8.13 | 2.01 | 0.522 | 0.122 | 0.048 |
| Average | 7.44 | 1.75 | 0.480 | 0.133 | 0.044 |
| SD | 0.680 | 0.142 | 0.038 | 0.007 | 0.003 |
| %CV | 9.1 | 8.1 | 7.9 | 5.1 | 6.7 |

**Table 3.11: Inter-assay Precision Data for the EBOV GP Monomer Serology Assay**

| Operator | Day | Plate | VHQC-1 (EU/ml) | HQC-1 (EU/ml) | MQC-1 (EU/ml) | LQC-1 (EU/ml) | VLQC-1 (EU/ml) |
|---|---|---|---|---|---|---|---|
| 1 | 1 | 1 | 8.32 | 2.10 | 0.538 | 0.155 | 0.085 |
| 1 | 2 | 1 | 10.5 | 2.33 | 0.575 | 0.190 | 0.116 |
| 1 | 2 | 2 | 8.14 | 2.08 | 0.536 | 0.178 | 0.101 |
| 1 | 3 | 1 | 8.59 | 2.51 | 0.611 | 0.170 | 0.077 |
| 1 | 4 | 1 | 9.16 | 2.00 | 0.551 | 0.184 | 0.103 |
| 1 | 4 | 2 | 8.42 | 2.37 | 0.573 | 0.174 | 0.096 |
| 1 | 5 | 1 | 8.46 | 2.00 | 0.516 | 0.154 | 0.076 |
| 1 | 6 | 1 | 9.32 | 2.27 | 0.541 | 0.185 | 0.102 |
| 2 | 7 | 1 | 8.99 | 1.88 | 0.503 | 0.197 | 0.128 |
| Average | | | 8.88 | 2.17 | 0.549 | 0.176 | 0.098 |
| SD | | | 0.734 | 0.208 | 0.033 | 0.015 | 0.017 |
| %CV | | | 8.3 | 9.6 | 6.0 | 8.4 | 17.7 |

**Table 3.12: Inter-assay Precision Data for the EBOV GP Monomer Serology Assay**

| Operator | Day | Plate | VHQC-2 (EU/ml) | HQC-2 (EU/ml) | MQC-2 (EU/ml) | LQC-2 (EU/ml) | VLQC-2 (EU/ml) |
|---|---|---|---|---|---|---|---|
| 1 | 1 | 1 | 8.32 | 2.17 | 0.558 | 0.148 | 0.049 |
| 1 | 2 | 1 | 9.24 | 2.04 | 0.574 | 0.177 | 0.062 |
| 1 | 2 | 2 | 8.21 | 1.99 | 0.497 | 0.152 | 0.055 |
| 1 | 3 | 1 | 7.95 | 2.22 | 0.540 | 0.138 | 0.039 |
| 1 | 4 | 1 | 8.17 | 2.15 | 0.565 | 0.165 | 0.057 |
| 1 | 4 | 2 | 7.57 | 2.23 | 0.556 | 0.154 | 0.055 |
| 1 | 5 | 1 | 7.44 | 1.75 | 0.480 | 0.133 | 0.044 |
| 1 | 6 | 1 | 8.67 | 2.08 | 0.509 | 0.162 | 0.070 |
| 2 | 7 | 1 | 7.26 | 1.98 | 0.574 | 0.182 | 0.086 |
| Average | | | 8.09 | 2.07 | 0.539 | 0.157 | 0.058 |
| SD | | | 0.626 | 0.151 | 0.035 | 0.016 | 0.014 |
| %CV | | | 7.7 | 7.3 | 6.6 | 10.5 | 24.7 |

**Table 3.13: Dilutional Linearity: Anti-EBOV GP Monomer Antibody Concentrations of Diluted Samples**

| Sample | Dilution Factor | GP Monomer (EU/ml) |
|---|---|---|
| PREVAIL 9 | 2,000 | 389 |
| | 4,000 | 347 |
| | 8,000 | 363 |
| | 16,000 | 346 |
| | 32,000 | 311 |
| PREVAIL 10 | 2,000 | 140 |
| | 4,000 | 171 |
| | 8,000 | 159 |
| | 16,000 | 128 |
| | 32,000 | 105 |
| PREVAIL 14 | 2,000 | 1,774 |
| | 4,000 | 1,748 |
| | 8,000 | 1,744 |
| | 16,000 | 2,014 |
| | 32,000 | 2,090 |
| PREVAIL 17 | 2,000 | 15,820 |
| | 4,000 | 18,260 |
| | 8,000 | 13,450 |
| | 16,000 | 14,280 |
| | 32,000 | 13,580 |
| PREVAIL 85 | 2,000 | 27,010 |
| | 4,000 | 27,570 |
| | 8,000 | 30,690 |
| | 16,000 | 37,500 |
| | 32,000 | 34,220 |

**Table 3.14: Dilutional Linearity of Diluted Samples**

| Sample | Dilution Factor | % Recovery 1:4,000 | % Recovery 1:8,000 |
|---|---|---|---|
| PREVAIL 9 | 2,000 | 112 | 107 |
| | 4,000 | 100 | 96 |
| | 8,000 | 105 | 100 |
| | 16,000 | 100 | 95 |
| | 32,000 | 90 | 86 |
| PREVAIL 10 | 2,000 | 82 | 88 |
| | 4,000 | 100 | 107 |
| | 8,000 | 93 | 100 |
| | 16,000 | 75 | 80 |
| | 32,000 | 62 | 66 |
| PREVAIL 14 | 2,000 | 101 | 102 |
| | 4,000 | 100 | 100 |
| | 8,000 | 100 | 100 |
| | 16,000 | 115 | 115 |
| | 32,000 | 120 | 120 |
| PREVAIL 17 | 2,000 | 87 | 118 |
| | 4,000 | 100 | 136 |
| | 8,000 | 74 | 100 |
| | 16,000 | 78 | 106 |
| | 32,000 | 74 | 101 |
| PREVAIL 85 | 2,000 | 98 | 88 |
| | 4,000 | 100 | 90 |
| | 8,000 | 111 | 100 |
| | 16,000 | 136 | 122 |
| | 32,000 | 124 | 112 |

**Table 3.15: Spike and Recovery: Anti-EBOV GP Monomer Antibody Concentrations of Spiked Samples***

| Sample | Spike | GP Monomer (EU/ml) |
|---|---|---|
| Prevail 2 | Very High | 11.5 |
| | High | 2.64 |
| | Medium | 0.734 |
| | Low | 0.185 |
| | Very Low | 0.043 |
| | None | **0.001** |
| Prevail 3 | Very High | 10.3 |
| | High | 2.74 |
| | Medium | 0.722 |
| | Low | 0.174 |
| | Very Low | 0.039 |
| | None | **0.0002** |
| Prevail 6 | Very High | 11.8 |
| | High | 2.73 |
| | Medium | 0.742 |
| | Low | 0.190 |
| | Very Low | 0.040 |
| | None | **0.0002** |
| Prevail 7 | Very High | 11.0 |
| | High | 2.49 |
| | Medium | 0.633 |
| | Low | 0.163 |
| | Very Low | 0.037 |
| | None | **0.0004** |
| Prevail 8 | Very High | 9.11 |
| | High | 2.34 |
| | Medium | 0.548 |
| | Low | 0.153 |
| | Very Low | 0.039 |
| | None | **0.0002** |

*Bold values <LLOQ

**Table 3.16: Spike and Recovery: Percent Recovery of Spiked Samples**

| Sample | Spike | % Recovery |
|---|---|---|
| Prevail 2 | Very High | 100 |
| | High | 87 |
| | Medium | 91 |
| | Low | 86 |
| | Very Low | 100 |
| Prevail 3 | Very High | 90 |
| | High | 90 |
| | Medium | 89 |
| | Low | 81 |
| | Very Low | 91 |
| Prevail 6 | Very High | 103 |
| | High | 90 |
| | Medium | 92 |
| | Low | 89 |
| | Very Low | 94 |
| Prevail 7 | Very High | 105 |
| | High | 88 |
| | Medium | 96 |
| | Low | 87 |
| | Very Low | 93 |
| Prevail 8 | Very High | 87 |
| | High | 83 |
| | Medium | 84 |
| | Low | 82 |
| | Very Low | 98 |

**Table 3.17: Spike and Recovery: Anti-EBOV GP Monomer Antibody Concentrations of Spiked Samples***

| Sample | Spike | GP Monomer (EU/ml) |
|---|---|---|
| Prevail 11 | Very High | 10.0 |
| | High | 2.38 |
| | Medium | 0.603 |
| | Low | 0.161 |
| | Very Low | 0.038 |
| | None | **0.001** |
| Prevail 12 | Very High | 10.4 |
| | High | 2.35 |
| | Medium | 0.577 |
| | Low | 0.153 |
| | Very Low | 0.037 |
| | None | **0.001** |
| Prevail 22 | Very High | 9.61 |
| | High | 2.47 |
| | Medium | 0.589 |
| | Low | 0.147 |
| | Very Low | 0.035 |
| | None | **0.001** |
| Prevail 25 | Very High | 10.8 |
| | High | 2.54 |
| | Medium | 0.647 |
| | Low | 0.169 |
| | Very Low | 0.037 |
| | None | **0.001** |
| Prevail 26 | Very High | 10.3 |
| | High | 2.05 |
| | Medium | 0.551 |
| | Low | 0.147 |
| | Very Low | 0.036 |
| | None | **0.002** |

*Bold values are <LLOQ

**Table 3.18: Spike and Recovery: Percent Recovery of Spiked Samples**

| Sample | Spike | % Recovery |
|---|---|---|
| Prevail 11 | Very High | 96 |
| | High | 84 |
| | Medium | 92 |
| | Low | 86 |
| | Very Low | 93 |
| Prevail 12 | Very High | 99 |
| | High | 85 |
| | Medium | 88 |
| | Low | 86 |
| | Very Low | 100 |
| Prevail 22 | Very High | 91 |
| | High | 89 |
| | Medium | 90 |
| | Low | 82 |
| | Very Low | 92 |
| Prevail 25 | Very High | 102 |
| | High | 92 |
| | Medium | 99 |
| | Low | 95 |
| | Very Low | 98 |
| Prevail 26 | Very High | 109 |
| | High | 93 |
| | Medium | 100 |
| | Low | 103 |
| | Very Low | 111 |

**Table 3.19: Concentration of anti-EBOV GP Monomer Antibodies in IRF Samples***

| Sample Name | Barcode | GP Monomer (EU/ml) |
|---|---|---|
| IRF 13 | 1-01374-3-02 | 42.2 |
| IRF 14 | 1-00608-3-02 | 1289 |
| IRF 15 | 1-01266-3-02 | 6967 |
| IRF 16 | 1-01225-3-02 | 159 |
| IRF 17 | 1-00325-3-02 | 10110 |
| IRF 18 | 1-01520-3-02 | 2098 |
| IRF 19 | 1-01597-3-02 | 1536 |
| IRF 20 | 1-01616-3-02 | 590 |
| IRF 21 | 1-01631-3-02 | 1036 |
| IRF 22 | 1-01593-3-02 | **13.7** |
| IRF 23 | 1-00275-3-02 | 380 |
| IRF 24 | 1-02512-3-02 | 2056 |
| IRF 25 | 1-01838-3-02 | **10.3** |
| IRF 26 | 1-02087-3-02 | **10.5** |
| IRF 27 | 1-02497-3-02 | 1348 |
| IRF 28 | 1-02893-3-02 | 3569 |
| IRF 29 | 1-02000-3-02 | 878 |
| IRF 30 | 1-00295-3-02 | **17.9** |
| IRF 31 | 1-02789-3-02 | 2093 |
| IRF 32 | 1-00259-3-02 | 2387 |
| IRF 33 | 1-02734-3-02 | 62.2 |
| IRF 34 | 1-02810-3-02 | 25.6 |
| IRF 35 | 1-03002-3-02 | 632 |
| IRF 36 | 3-00158-4-02 | 703 |
| IRF 37 | 3-00166-4-02 | **12.2** |
| IRF 38 | 3-00325-4-02 | 7302 |
| IRF 39 | 3-00342-4-02 | 2457 |
| IRF 40 | 3-00366-4-02 | 233 |
| IRF 41 | 3-00356-4-03 | 1430 |
| IRF 45 | 3-00677-4-03 | 1996 |

*Bold values are <LLOQ

**Table 3.20: Concentration of anti-EBOV GP Monomer Antibodies in IRF Samples***

| Sample Name | Barcode | GP Monomer (EU/ml) |
|---|---|---|
| IRF 61 | 3955302 | **11.6** |
| IRF 62 | 4258302 | 45.3 |
| IRF 63 | 1891302 | 3086 |
| IRF 64 | 1788302 | 10290 |
| IRF 65 | 2078302 | 16560 |
| IRF 66 | 2132302 | 34.7 |
| IRF 67 | 2197302 | 86450 |
| IRF 68 | 2236302 | 7764 |
| IRF 69 | 2310302 | 5666 |
| IRF 70 | 90070302 | 24130 |
| IRF 71 | 2687302 | **15.4** |
| IRF 72 | 2769302 | 271 |
| IRF 73 | 3079302 | 6707 |
| IRF 74 | 3085302 | **12.1** |
| IRF 75 | 3263302 | 43.9 |
| IRF 76 | 3561302 | 6123 |
| IRF 77 | 3805302 | 2211 |
| IRF 78 | 4296302 | 1413 |
| IRF 79 | 128848301 | 1173 |
| IRF 80 | 129521301 | 3224 |
| IRF 81 | 129207301 | 2849 |
| IRF 82 | 129830301 | 53.3 |
| IRF 83 | 129742301 | 3705 |
| IRF 85 | 129026301 | 32380 |
| IRF 86 | 129046301 | 4099 |
| IRF 87 | 128891301 | 10140 |
| IRF 88 | 129380301 | 2360 |
| IRF 91 | 129157301 | 1098 |
| IRF 93 | 129258301 | 1315 |
| IRF 95 | 129691301 | 1930 |

*Bold values are <LLOQ
